# Adaptive remodeling of rat adrenomedullary stimulus-secretion coupling in response to a chronic hypertensive environment

**DOI:** 10.1101/2023.11.28.568973

**Authors:** Vincent Paillé, Joohee Park, Bertrand Toutain, Jennifer Bourreau, Pierre Fontanaud, Frédéric De Nardi, Claudie Gabillard-Lefort, Dimitri Bréard, David Guilet, Daniel Henrion, Christian Legros, Nathalie C. Guérineau

**Author notes:** To whom correspondence should be addressed at Institut de Génomique Fonctionnelle, Université Montpellier; CNRS UMR5203; INSERM U1191; 141 rue de la Cardonille, 34094 Montpellier CEDEX 05, France.

## Abstract

Chronic elevated blood pressure impinges on the functioning of multiple organs and therefore harms body homeostasis. Elucidating the protective mechanisms whereby the organism copes with sustained or repetitive blood pressure rises is therefore a topical challenge. Here we address this issue in the adrenal medulla, the master neuroendocrine tissue involved in the secretion of catecholamines, influential hormones in blood pressure regulation. Using acute adrenal slices from spontaneously hypertensive rats, we show that chromaffin cell stimulus-secretion coupling is remodeled, resulting in a less efficient secretory function primarily upon sustained electrical or cholinergic challenges. The remodeling is supported by revamped cellular and tissular mechanisms, including chromaffin cell excitability through voltage-gated ion channel expression changes, gap junctional communication and cholinergic synaptic transmission. As such, by weakening its competence to release catecholamines, the ‘hypertensive medulla’ has elaborated an adaptive shielding mechanism against damaging effects of redundant elevated catecholamine secretion and associated blood pressure.

## Introduction

A rise in circulating catecholamine (CA) levels is a crucial step triggered by the organism to cope with a stressful situation. By releasing both epinephrine (E) and norepinephrine (NE), the adrenal medullary tissue crucially contributes to this response. Beyond the beneficial effect of CA secretion elicited by an acute stress, sustained and/or repetitive CA secretion episodes (in response to chronic stressful situations for example) can have deleterious outcomes ^1^, as unveiled by the elevated blood pressure observed in response to chronic infusion of E in rat ^2,3^ or in chronically cold stressed rats ^4^. It has long been reported that increased blood pressure is associated with a sympathetic nervous system hyperactivity leading to a raised neural tone ^5,6^ and increased plasma E and NE ^7^. The adrenal medulla being the unique source of circulating E, this assigns the secretory function of the medulla and more generally the sympatho-adrenal axis as critical determinants of arterial hypertension pathogenesis ^8–10^. Reciprocally, and despite it is indisputable that the adrenal medulla primarily contributes to enhance blood pressure, whether and how a ‘hypertensive’ environment impacts the adrenal secretory function itself remains elusive. We address this issue in the adult spontaneously hypertensive rat, in which the circulating E levels gradually increase as arterial hypertension develops and then stabilize in adult animals to become comparable to those in normotensive rats ^11^. gis suggests that the adrenal gland develops shielding mechanisms dedicated to normalize blood CA amounts, and to date nothing is known on the adaptive mechanisms elaborated by the neuroendocrine chromaffin cells to adjust the release of CA.

Adrenal CA secretion and more generally the adrenomedullary tissue function are controlled by the coordination of interconnected and complex pathways ^12^. The initial incoming command comes from the sympathetic nervous system that releases mainly acetylcholine (ACh) but also neuropeptides (PACAP, VIP…) at splanchnic nerve terminals synapsing onto chromaffin cells ^13,14^. The resulting ACh-evoked chromaffin cell depolarization and subsequent cytosolic Ca^2+^ rise are key processes for CA exocytosis. In addition and unraveled from studies in both acute adrenal slices and *in vivo* in anaesthetized rodents, the local communication mediated by gap junctions between chromaffin cells represents a functional route by which biological signals propagate between adjacent cells and subsequently contribute to CA release^15–20^.

To elucidate the feedback mechanisms elaborated by the adrenal medullary tissue to manage a chronic sustained elevated blood pressure, we investigated chromaffin cell stimulus-secretion coupling in acute adrenal slices of adult spontaneously hypertensive rats (SHRs) and their parent age-matched normotensive Wistar Kyoto (WKY) rats. By driving numerous cellular/tissular processes, chromaffin cell excitability is a major player in stimulus-secretion coupling. It relies on intricate mechanisms, not only supported by ion channels expressed at the plasma membrane, but also by the crosstalk between cholinergic and peptidergic innervation ^21^ and the gap junctional electrical coupling between chromaffin cells ^17,19,20^. We identified relevant modifications in chromaffin cell excitability through changes in voltage-gated ion channel expression, cholinergic synaptic neurotransmission and gap junctional communication, that argue for a less efficient stimulus-secretion coupling in hypertensive animals, markedly observed upon robust challenges. We propose that this functional plasticity reflects an adaptive shielding mechanism, avoiding the detrimental effects of sustained or repetitive huge CA secretion episodes. More generally, this study describes novel adaptive mechanisms that take place in the medullary tissue and how they act in a coordinated manner to damper CA release.

## Results

### Impaired CA release evoked by high ACh concentration in hypertensive rats

The competence of WKY and SHR chromaffin cells to secrete CA was investigated by assaying basal and ACh-evoked epinephrine (E) and norepinephrine (NE) released from acute slices (Figures 1A, 1B and 1C). We took advantage of making slices from the two adrenals to estimate the medulla volume of the right and left glands for both SHRs and WKY rats, as previously reported ^22^. For each rat, whether WKY or SHR, no difference was observed between the right and the left glands (0.85 ± 0.08 mm^3^, n = 4 and 0.95 ± 0.15 mm^3^, n = 4 for the right and left WKY glands, respectively, p = 0.125, and 1.17 ± 0.33 mm^3^, n = 4 and 1.25 ± 0.32 mm^3^, n = 4 for the right and left SHR glands, respectively, p = 0.125, Wilcoxon matched-pairs signed-rank test, Figure S1D). However, medullary tissue volume was significantly greater in SHRs (1.21 ± 0.30 mm^3^, n = 8 glands for SHRs *versus* 0.90 ± 0.12 mm^3^, n = 8 glands for WKY rats, p = 0.0351, Mann-Whitney test, Figure S1E). To make appropriate comparisons between SHRs and WKY rats, CA secretion was normalized per mm^3^ medulla, as previously described ^22^. Under basal conditions, SHR chromaffin cells secreted 8.6-fold less NE than WKY cells (0.7 ± 0.7 pmol/mm3, n = 11 slices for SHRs *versus* 5.9 ± 7.4 pmol/mm3, n = 10 slices for WKY rats, p = 0.0003, Mann-Whitney test, Figure 1A, right histogram). No change was observed for released E amounts (40.3 ± 22.6 pmol/mm3, n = 11 slices for SHRs *versus* 40.0 ± 18.3 pmol/mm3, n = 9 slices for WKY rats, p = 0.766, Mann-Whitney test, Figure 1A, left histogram). Likewise, a significant decrease in NE release in SHRs was observed in response to ACh stimulations, both at low (1 μM) and high (10-100 μM) concentrations (Figure 1B, p = 0.0018, 0.0015 and 0.0002 for ACh 1, 10 and 100 μM, respectively, Mann-Whitney test). The E release amount is also significantly decreased, but only at 100 μM ACh (p = 0.2945, 0.5269 and 0.0147 for ACh 1, 10 and 100 μM, respectively, Mann-Whitney test). To go further, we investigated the ability of chromaffin cells to be stimulated by ACh. Irrespective to ACh concentrations (1-100 μM), the stimulation ratio was similar between SHRs and WKY rats, even at high ACh concentration (Figure 1C, for E: x5.9 ± 3.5 (n = 15 slices) and x4.0 ± 1.5 (n = 11 slices) for WKY rats and SHRs, respectively, p = 0.1447, Mann-Whitney test and for NE: x4.9 ± 2.9 (n = 15 slices) and x5.3 ± 2.1 (n = 12 slices) for WKY rats and SHRs, respectively, p = 0.3406, Mann-Whitney test). Strengthening a deficiency in ACh-induced E secretion in SHRs, the E:NE ratio remained constant upon challenging SHR chromaffin cells with increasing ACh concentrations (p = 0.7061, n = 4 rats, Kruskal-Wallis test), while it gradually increased in WKY rats, as expected (Figure 1D). Altogether, this indicates that despite a preserved ability to release CA, stimulus-secretion coupling of chromaffin cells is less efficient in SHRs. Aiming at identifying which element(s) of the adrenal stimulus-secretion is (are) altered in SHRs, we next performed a series of experiments focused on i) chromaffin cell excitability, ii) the cholinergic neurotransmission at the splanchnic nerve ending-chromaffin cell synapse and iii) the gap junction-mediated communication between chromaffin cells.

**Figure 1:**
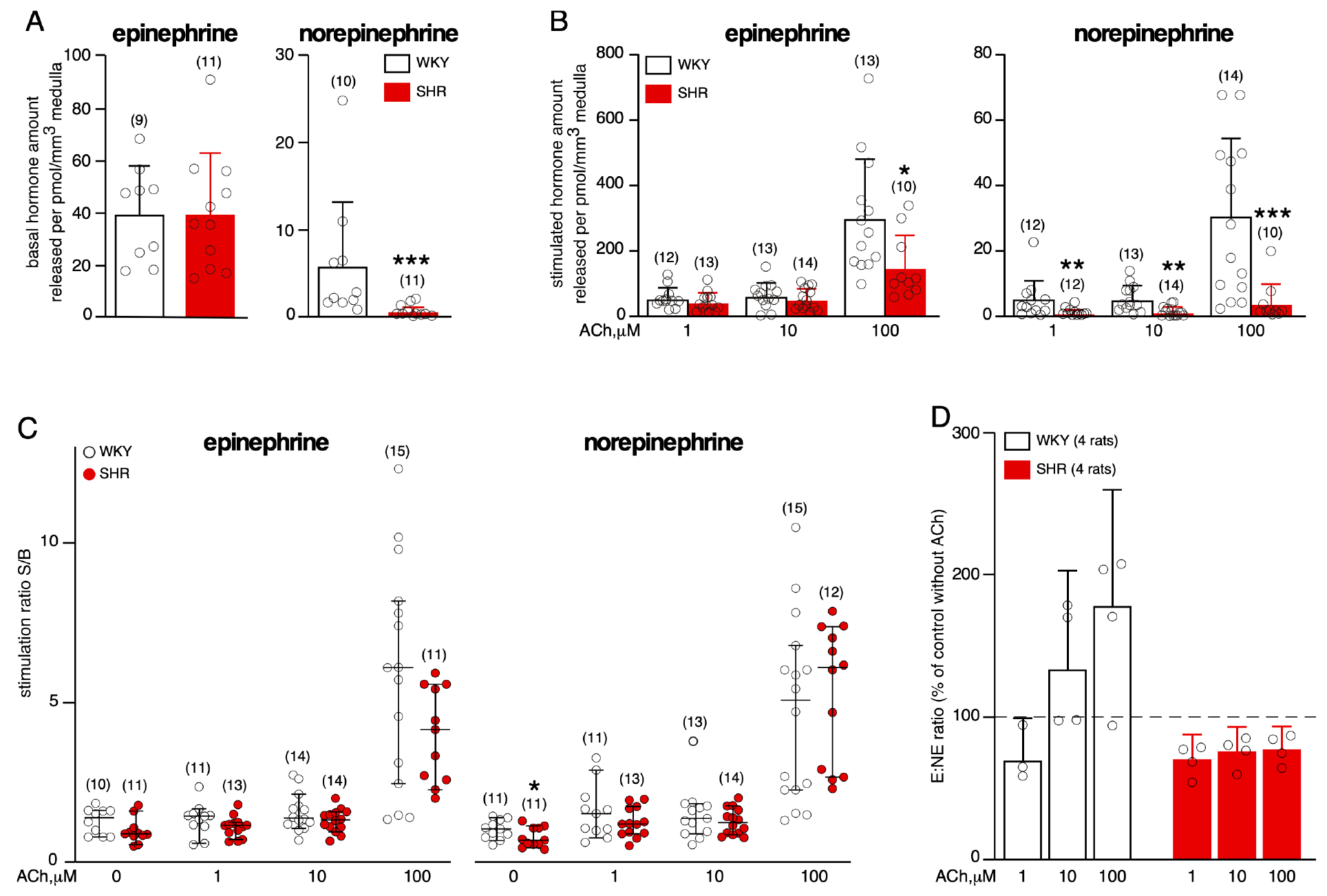
Reduced competence to release CA in SHRs in response to robust ACh challenge. Acute adrenal slices from SHRs and WKY rats were incubated first for 5 min (basal (B) conditions) and then challenged (stimulated (S) conditions) with either ACh-containing saline or Ringer saline, during 5 min. Secreted epinephrine (E) and norepinephrine (NE) were assayed by HPLC. **A.** Basal CA secretion. **B.** CA secretion in response to increasing ACh stimulations. Both basal and ACh-evoked NE amounts released per mm^3^ of medulla are significantly reduced in SHRs. Regarding E, amounts released per mm^3^ are decreased only in response to a high ACh concentration (100 μM)-evoked challenge. **C.** No difference in stimulation ratios S/B, indicating that the tissue responsiveness is preserved in SHRs. The number of slices is indicated in parentheses. **D.** E:NE ratios calculated from the two adrenals pooled for each rat. Constant E:NE ratios in SHRs, contrasting with a progressive increase in E:NE in WKY rats, upon exposure to increasing concentrations of ACh.

### Altered action potential firing in SHRs in response to sustained stimulations

Regarding the passive electrical membrane properties of chromaffin cells, significant changes were observed between SHRs and WKY rats for input resistance and capacitance (Table S1), predicting a plausible stimulus-secretion coupling reshaping in SHRs. To investigate chromaffin cell excitability under experimental conditions as close to *in situ* conditions as possible, spontaneous action potential (AP) firing at resting membrane potential were recorded in the loose cell-attached configuration (Figure 2A). Neither the percentage of chromaffin cells exhibiting spontaneous APs (80.5 %, n = 33/41 cells and 81.6%, n = 31/38 cells for WKY and SHRs, respectively, p>0.9999, Fisher’s exact test) nor the discharge frequency (0.69 ± 1.07 Hz, n = 32 cells for WKY rats and 0.45 ± 0.63 Hz, n = 31 cells for SHRs, p = 0.2964, unpaired t test, Figure 2B, left histogram) significantly differed between SHRs and WKY rats. Likewise, the frequency distribution was not different between SHRs and WKY rats, p>0.9999, Fisher’s exact test, Figure 2B, right histogram).

**Figure 2:**
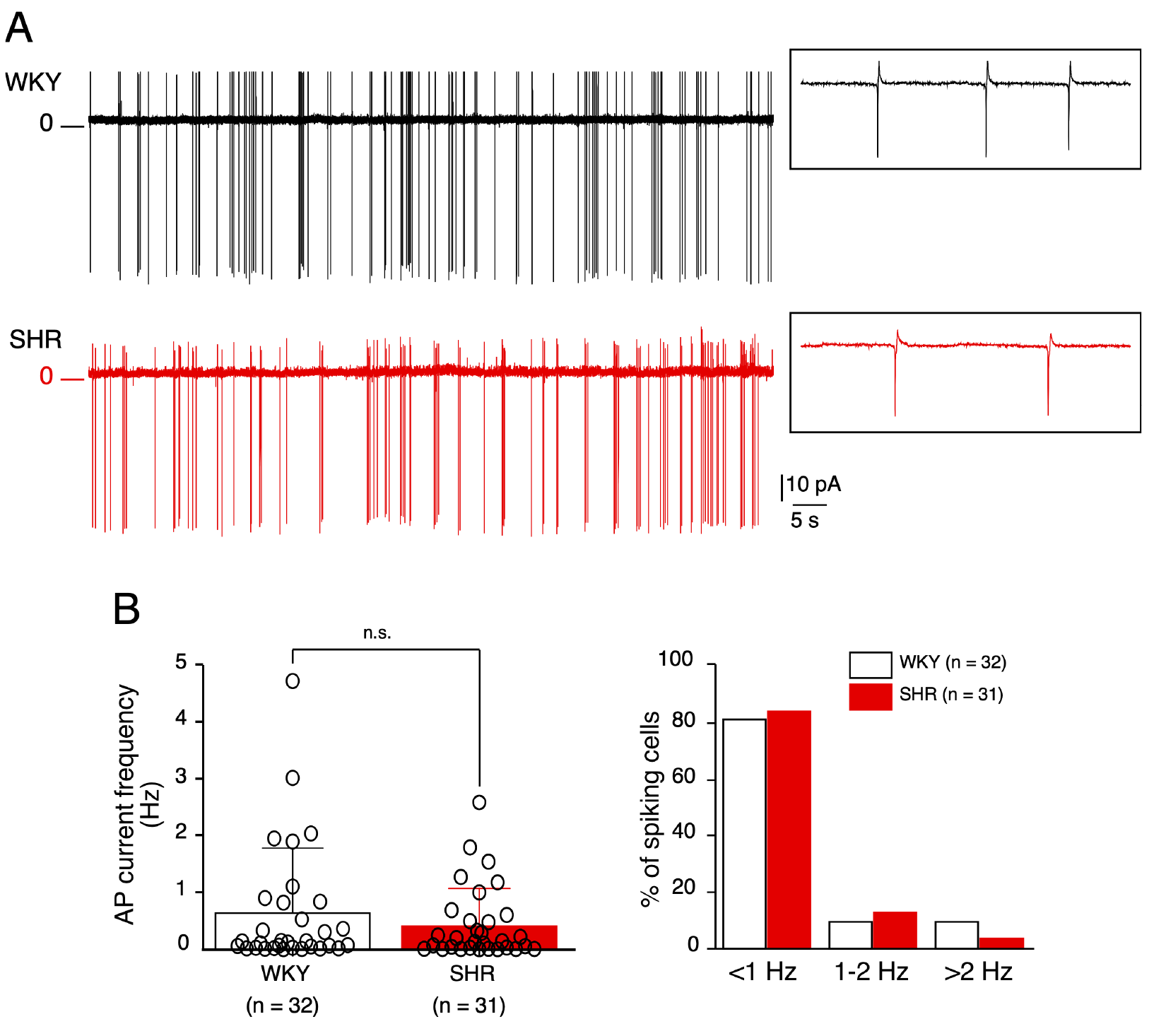
Spontaneous electrical firing monitored in chromaffin cells in adrenal acute slices of WKY rats and SHRs. Action potential (AP) currents were recorded in the voltage-clamp mode (0 mV) of the loose cell-attached configuration. **A.** Representative chart recordings of a WKY (upper trace) cells and a SHR (lower trace) cell. **B.** Analysis of the AP current frequency. Neither the mean frequency nor the distribution (ranging from <1 to >2 Hz) differed between WKY rats and SHRs.

The loose patch configuration allowing for long-lasting recordings, we next analyzed the spiking pattern of WKY and SHR chromaffin cells. Do they exhibit a regular and/or a bursting pattern, as previously identified in mouse chromaffin cells ^23^? To discriminate between the two firing patterns, the regularity of the firing discharge was investigated by calculating the coefficient of variation (CV) of inter-spike interval (ISI) distribution. The analysis of the discharge pattern (150-200 s recording) was performed on 24 WKY and 26 SHR chromaffin cells, and shows that WKY and SHR cells both fire regularly or in bursts (Figures 3A and 3B and Figure S2). As expected and consistent with previous data in mouse chromaffin cells ^23^, a regular firing mode is associated with a CV<1 and a bursting mode with a CV>1. No overt difference in the percentage of cells displaying a regular or a bursting firing was observed between SHR and WKY rats (3 regular and 23 bursting cells (n = 26) for SHR and 4 regular and 20 bursting cells (n = 24) for WKY, respectively, p = 0.6971, Fisher’s exact test, Figure 3C). To determine whether the two firing patterns account for two distinct chromaffin cell populations or can occur in the same cell, the spontaneous electrical activity of individual chromaffin cells was recorded for a longer period of time (5-15 minutes, loose-patch configuration). Interestingly, and both in WKY and SHR rats, the firing discharge can alternate between the two modes, switching from a regular to a bursting mode and vice-versa (Figure S3). This clearly indicates that these spiking patterns do not reflect two different cell populations.

**Figure 3:**
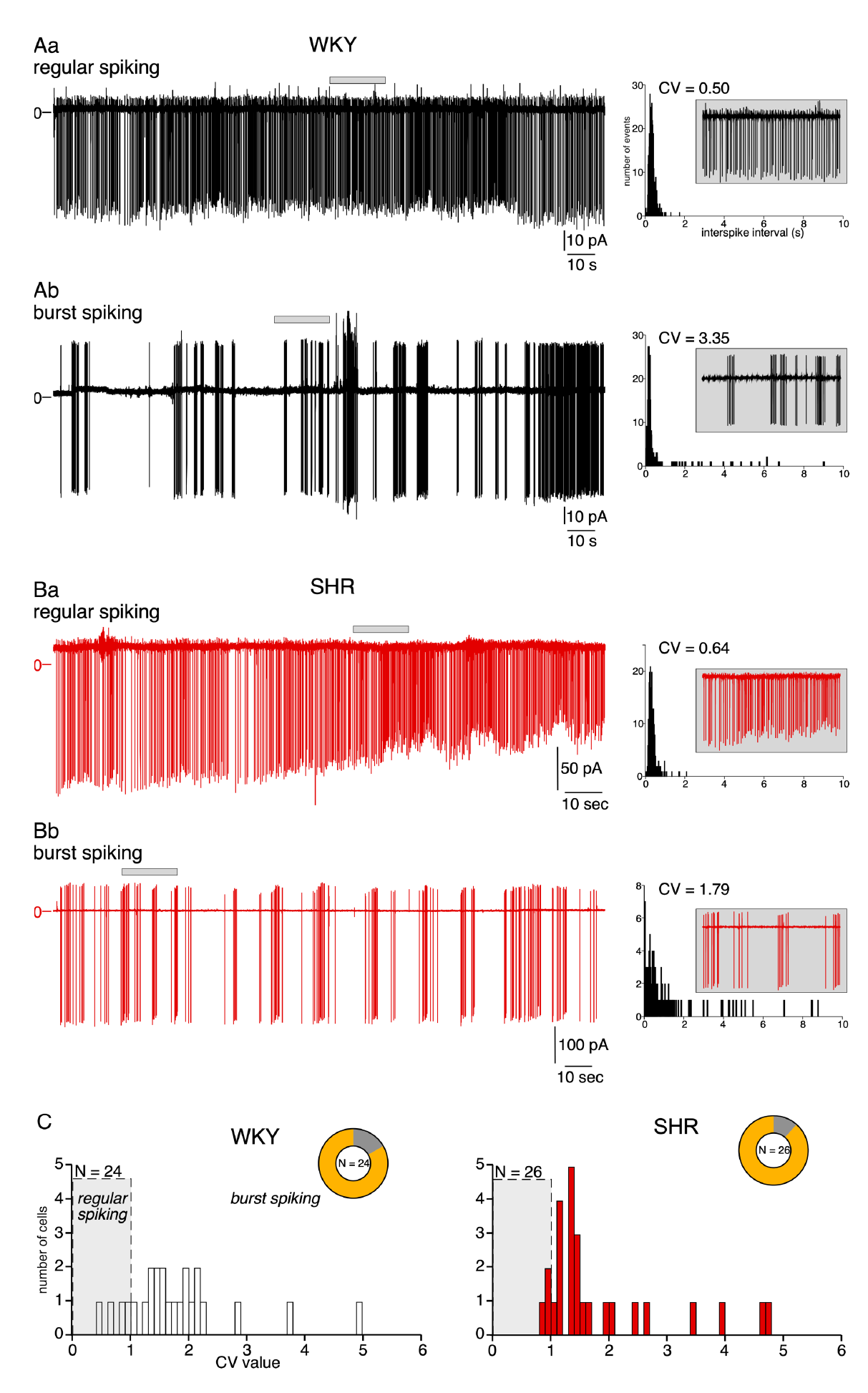
Presence of two distinct firing patterns in chromaffin cells of WKY rats and SHRs. Cells were recorded in the loose cell-attached configuration and voltage-clamped at 0 mV. **A.** Spontaneous AP currents recorded in two individual cells of WKY rats. One cell exhibits a regular spiking (**Aa**, coefficient of variation CV = 0.5), while the second cell displays a bursting firing pattern (**Ab**, CV = 3.35). The histograms on the right illustrate the distribution of the inter-spike intervals (100 ms bin), from which the coefficients of variation (CV) were calculated, as described in ^23^. **B.** Spontaneous AP currents recorded in two individual cells of SHRs. One cell exhibits a regular spiking (**Ba**, coefficient of variation CV = 0.64), while the second cell displays a bursting firing pattern (**Bb**, CV = 1.79). Insets in A and B: expanded time scale illustrating a 20-s spiking period. **C.** Histograms illustrating the distribution of the mean CV values calculated in 24 WKY cells and 26 SHR cells (0.1 CV unit bin).

To further investigate chromaffin cell excitability, depolarizing steps were evoked using the whole-cell configuration in cells current-clamped at their resting membrane potential (between -60 and -70 mV) (Figure 4A). APs were elicited during 500 ms from a rheobase of 14.8 ± 9.4 pA (n = 26 cells) for WKY rats and 11.5 ± 5.7 pA (n = 24 cells) in SHRs (p = 0.1479, unpaired t test). When plotting the AP frequency as a function of the injected current intensity, no significant difference was found between SHRs and WKY rats in response to small depolarizing currents (≤40 pA). Conversely, when injecting large current amplitudes (>40 pA), the AP accommodation increased in SHRs, leading to a significant reduction in firing frequency at 60 pA (28.5 ± 4.4 Hz, n = 13 for WKY rats *versus* 23.0 ± 5.0 Hz, n = 12 for SHRs, p = 0.0087, Mann-Whitney test, Figure 4B). gis was associated with a gradual drop of the AP peak (−34% between the second AP and the last AP in SHRs *versus* -25% in WKY rats) and a spike widening (+20% between the second AP and the last AP in SHRs *versus* +16% in WKY rats, p<0.001, Figure 4C). gis observation is particularly relevant in that robust depolarizations may simulate the elevated peripheral sympathetic tone associated with elevated blood pressure. For spontaneous APs recorded in current-clamp mode without current injection, as for evoked APs, the duration was higher (+63% half-width) and the amplitude was lower in SHRs compared to WKY rats (-11%, Figure 4D, mean of 30-50 successive APs). The overall features of evoked and spontaneous APs in SHRs and WKY rats are summarized in Table S2. Collectively, these results indicate that in response to sustained electrical stimulations, chromaffin cells of SHRs are less excitable than those of WKY rats.

**Figure 4:**
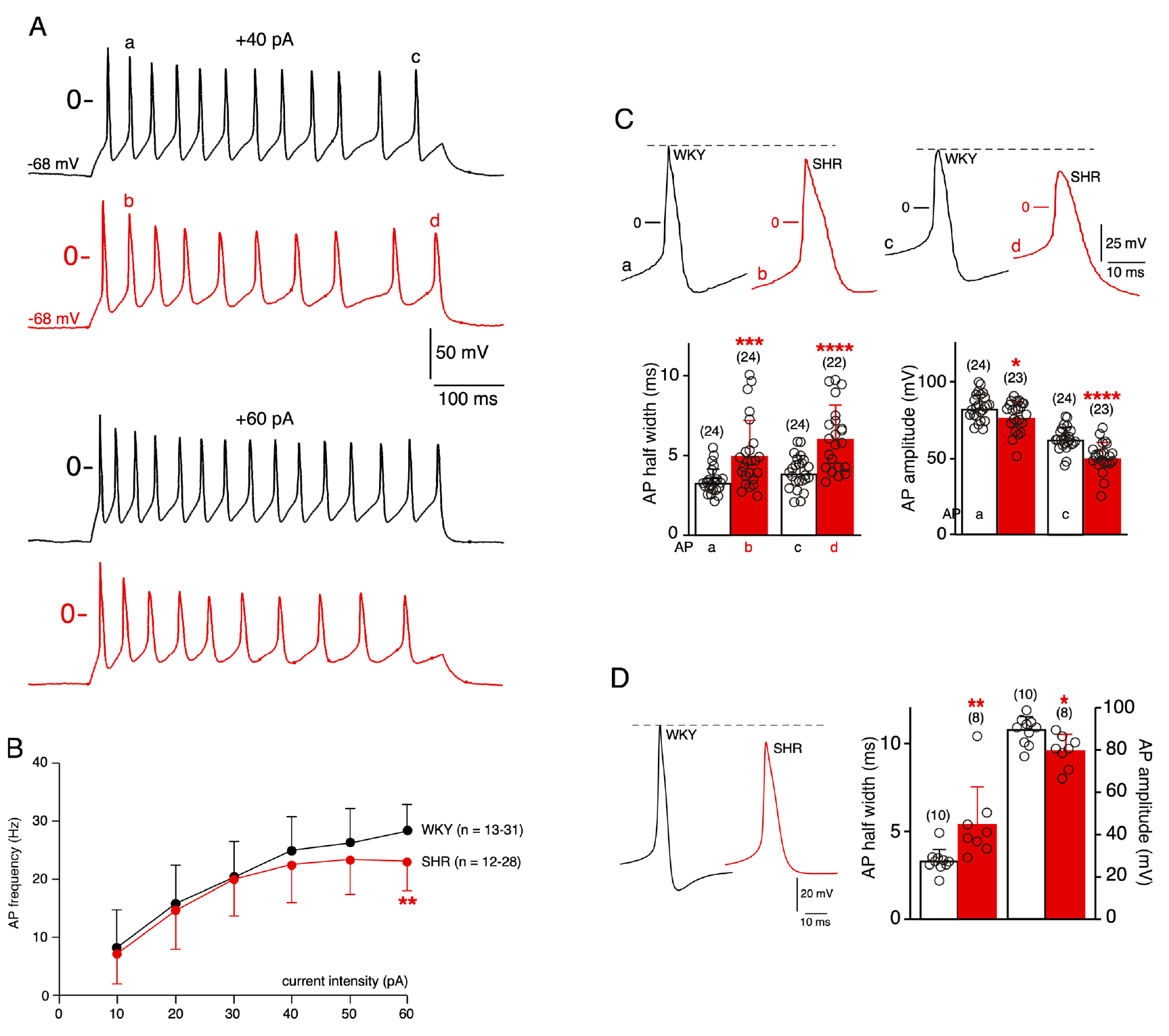
Decreased chromaffin cell excitability in response to robust depolarizations and AP waveform changes in SHRs. **A.** Illustrative chart recordings of evoked electrical activities in current-clamped WKY (black traces) and SHR (red traces) cells, in response to depolarizing steps (500 ms duration, +40 and +60 pA injected currents as upper and bottom traces, respectively). **B.** Analysis of the AP frequency in response to serial increasing depolarizations. Robust depolarizing steps (Iinj ≥40 pA) elicited less APs in SHRs that in WKY rats. **C.** Single AP waveforms derived from the second (a and b traces for WKY and SHR, respectively) and the last (c and d traces for WKY and SHR, respectively) evoked APs in response to a +40 pA, 500 ms duration depolarizing step. Analysis of AP half-width and amplitude changes shows an increased half-width and a decreased amplitude in SHRs. **D.** Similar analysis performed for spontaneous APs (mean of 30-50 APs/cell) recorded in WKY and SHR chromaffin cells current-clamped at their resting membrane potential. AP half width and amplitude are also significantly modified in SHRs (increased and decreased, respectively).

Does a decreased cell excitability occur in response to physiological stimuli? To address this issue, we studied the electrical behaviour of SHR and WKY chromaffin cells in response to the pituitary adenylyl cyclase-activating polypeptide (PACAP), the major neurotransmitter for stress transduction at the adrenomedullary synapse ^24,25^. Cells were recorded in the loose-patch cell-attached configuration and stimulated with 0.1 to 10 μM PACAP (Figure 5). Representative chart recordings in response to 10 μM PACAP are plotted in Figure 5A. AP currents were monitored before and after PACAP stimulation and the results were expressed as a percentage increase in AP frequency after PACAP (Figures 5B and 5C). PACAP, at 1 and 10 μM, increased the frequency of AP currents in WKY chromaffin cells, but not in SHRs. It should be noted that at higher PACAP concentration (10 μM), AP current frequency is even decreased when compared to that observed before PACAP application. gis result corroborates the decrease in excitability reported in Figure 4. To investigate whether the change in cell excitability reflects changes in the firing (regular *versus* bursting) pattern, we analysed the coefficient of variation, before and after PACAP exposure (0.1 to 10 μM, Figure S4). Although there is a downward trend, no significant differences were observed in both SHRs and WKY rats (2.22 ± 1.20 before PACAP and 1.89 ± 0.91 after PACAP, n = 12 cells for WKY rats, p = 0.2661, Wilcoxon matched-pairs signed-rank test and 2.25 ± 1.26 before PACAP and 1.62 ± 0.50 after PACAP, n = 10 cells, p = 0.1934, Wilcoxon matched-pairs signed-rank test, Figure S4A). In response to PACAP, the CV variance decreased more markedly in hypertensive than in normotensive animals, suggesting that PACAP acts by homogenizing the firing pattern of SHR chromaffin cells (Figure S4B). gis is consistent with the lack of correlation observed in SHRs, between CV after and before PACAP, in contrast to WKY rats (ρ = 0.713 in WKY rats, p = 0.0012, n = 12 cells, Spearman’s rank correlation coefficient *versus* ρ = 0.038 in SHRs, p = 0.916, n = 10 cells, Spearman’s rank correlation coefficient, Figures S4Ca and S4Cb). Altogether, these data obtained in response to PACAP, the main neurotransmitter operating at the ’stressed’ splanchnic nerve-chromaffin cell synapse, open up a new avenue of investigation. From a physiological point of view, the reduced ability of chromaffin cells to be stimulated by PACAP suggests an alteration of the stress response in SHRs, likely at the catecholamine secretion level.

**Figure 5:**
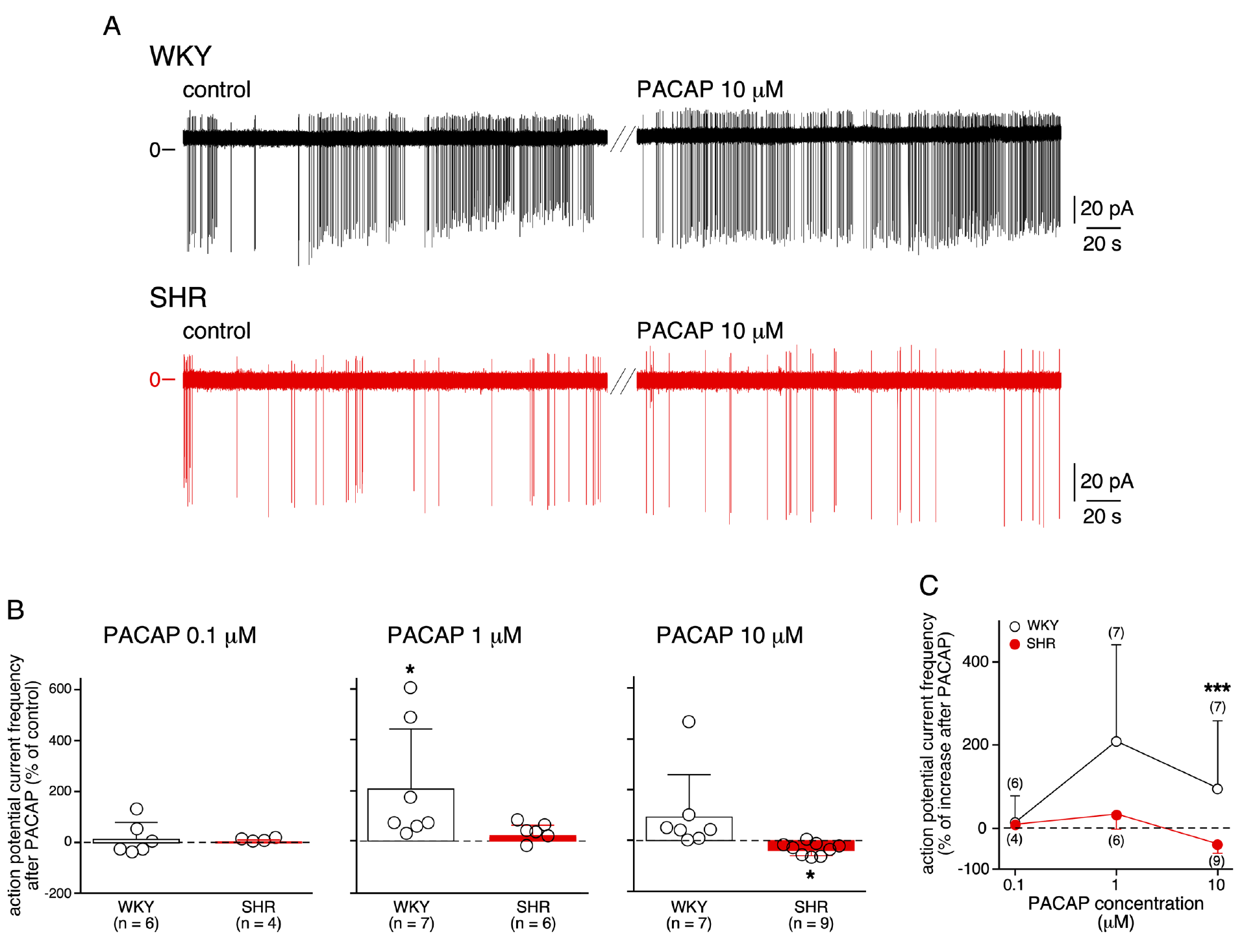
Physiological consequence of reduced excitability in SHRs: lack of electrical response to PACAP. Chromaffin cells in acute adrenal slices were recorded in the loose-patch cell-attached configuration (clamped at 0 mV) and stimulated with 0.1 to 10 μM PACAP-38. **A.** Representative chart recordings of AP currents in a WKY and SHR chromaffin cell exposed to 10 μM PACAP. **B.** Pooled data of electrical responses to various PACAP concentration applications, illustrating the lack of response to PACAP in SHRs compared to the increase in frequency of AP currents in WKY rats. The data are presented as a percentage relative to the firing frequency observed under control conditions. Note the decreased frequency in hypertensive animals upon a challenge with a high PACAP concentration (10 μM). **C**. Plot summarizing the effects of PACAP and the difference between SHRs and WKY rats, in particular at high PACAP concentration.

AP waveform critically relies on the nature and the relative proportion of voltage-gated ion channels. To identify the molecular determinants that could underlie the changes in AP shape, we investigated the expression of transcripts encoding voltage-dependent ion channels by quantitative PCR (Figure 6). We first investigated the expression level of mRNA encoding the α and auxiliary β subunits of the voltage-gated sodium (NaV1) channel family. As previously reported in mouse ^26^, *Scn3a* gene (encoding NaV1.3) is the main NaV channel gene expressed in the adrenal medulla of both normotensive and hypertensive rats (Figures 6A and 6C), with a significant upregulation in SHRs at the transcript level (1.4-fold increase, p = 0.0260, Mann-Whitney test, Figure 6B), which was not observed at the protein level (Figure S5). Regarding the other NaV1 isoforms, it is noteworthy that the expression level of *Scn9a* mRNA (encoding NaV1.7) displayed a 11.5-fold increase in SHRs (p = 0.0049, Mann-Whitney test, Figure 6B), accompagnied by a 1.3-fold increase at the protein level (p = 0.0411, Mann-Whitney test, Figure S5). NaV1.1, NaV1.4 and NaV1.5 transcripts (*Scn1a*, *Scn4a* and *Scn5a* genes, respectively) were not detected. Regarding NaV1 channel β subunits, *Scn1b* (NaVβ1 subunit), *Scn2b* (NaVβ2 subunit) and *Scn3b* (NaVβ3 subunit) genes were amplified, but not *Scn4b*. A significant increase was observed for *Scn2b* (2.5-fold change, p = 0.0022, Mann-Whitney test, Figure S6). With respect to Ca^2+^ channels, we examined both high threshold-activated CaV1 (L-type) and CaV2 (N-, P/Q- and R-type) channels, highly expressed in chromaffin cells and low threshold-activated CaV3 (T-type), whose expression undergoes robust remodeling in response to stressful situations ^27,28^. *Cacna1d* (encoding CaV1.3) and *Cacna1b* (encoding CaV2.2) genes are the principal transcripts expressed in WKY rats and SHRs, with respectively a 2-fold and a 1.5-fold increase in hypertensive animals (p = 0.0022 for *Cacna1d* and p = 0.041 for *Cacna1b*, Mann-Whitney tests, Figures 6A and 6B). The highest expression change was found for CaV2.1 channels (2.7-fold increase in SHRs, p = 0.0022, Mann-Whitney test). Transcripts encoding CaV3.2 (*Cacna1h*) and CaV3.3 channels (*Cacna1i*) were weakly expressed and their expression level did not change between normotensive and hypertensive rats. *Cacna1g* mRNA (encoding CaV3.1) was not detected. Concerning K^+^channel genes, we focused on Ca^2+^-activated channels expressed in chromaffin cells ^29^, namely the large (BK) KCa1.1 (*Kcnma1* gene), the intermediate (IK) KCa3.1 (*Kcnn4*) and the small (SK) KCa2 conductance channels (*Kcnn1-3*). Still, the overall collection of channels expressed (with a prevalent expression for KCa1.1 and KCa2.3) does not qualitatively differ between WKY and SHRs (Figure 6A). Quantitatively, the expression level of all transcripts, except *Kcnn2*, was significantly enhanced in SHRs (1.9-fold change for KCa1.1, 1.6-fold change for KCa2.1, 1.5-fold change for KCa2.3 and 1.8-fold change for KCa3.1). KCa4.1 (*Kcnt1*) and KCa4.2 (*Kcnt2*) mRNA were not detected. Altogether, these results show that i) the overall repertoire of voltage-gated ion channels expressed in WKY and SHR chromaffin cells is qualitatively similar (at the transcript level), except for the NaV1 channel family for which NaV1.7 (*Scn9a*) channels become the second most expressed isoform in SHRs after NaV1.3 channels and ii) the expression ratio between WKY and SHR channels for a given channel undergoes remodeling (Figure 6C).

**Figure 6:**
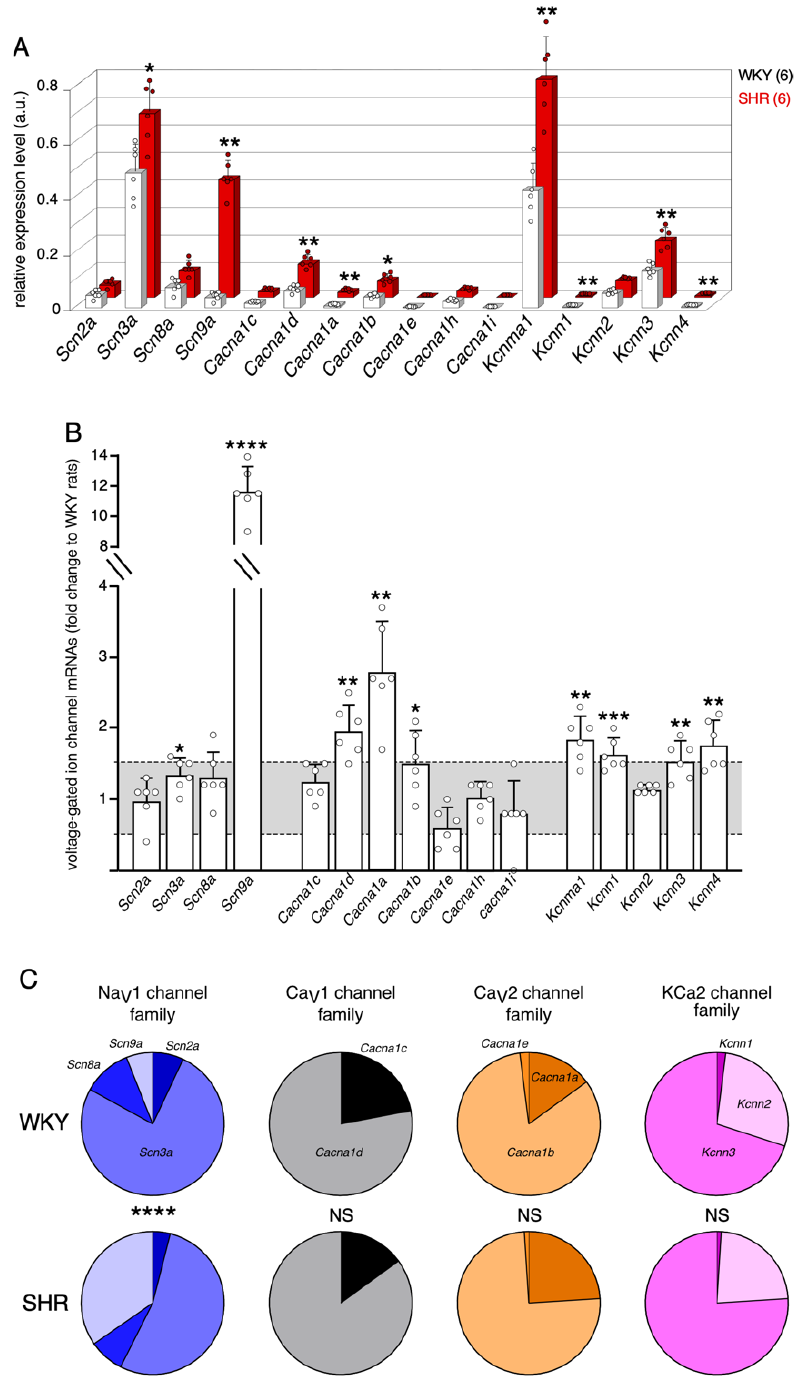
Remodeling of transcripts encoding voltage-gated ion channels in SHRs. Changes in mRNA expression levels were assessed by real-time RT-PCR in macrodissected adrenal medullary tissues from 6 WKY rats and 6 SHRs. **A.** 3D-histograms illustrating the relative expression levels in NaV, CaV and KCa channel genes. **B.** Fold changes in SHRs, as compared to WKY rat. The highest fold-change refers to *Scn9a* (encoding NaV1.7, 11.7-fold increase) for the NaV family, *Cacna4a* (encoding CaV2.1, >2.8-fold increase) for the CaV gene family and *Kcnma1* (encoding KCa1.1, 1.9-fold increase) for the KCa gene family. Fold change values were determined according to Livak’s method ^85^. The Shapiro-Wilk test was used to analyze the normality of data distribution, and parametric or non-parametric unpaired tests were used when appropriate. Fold changes between x0.5 and x1.5 (grey area) are considered irrelevant. **C.** Distribution of the channel isoforms in WKY rats and SHRs showing that the same voltage-gated channel families are present in the two strains, but with different expression ratios. Note the significant change for transcripts encoding NaV1 channel family, associated with a decrease for *Scn3a* and an increase for *Scn9a*.

Are those expression changes in voltage-gated ion channels relevant for cell electrical firing and can they account for the reduced chromaffin cell excitability that occur in SHRs in response to sustained depolarizations? To address this issue, the experimentally observed changes in the expression of Na^+^, Ca^2+^ and Ca^2+^-dependent K^+^ channels were evaluated in a numerical model of the rat chromaffin cell electrical activity ^30^ (Figure 7). The ‘WKY’ and ‘SHR’ models were implemented by the respective experimental values of resting membrane potential and membrane capacitance (see Table S1). For the ‘WKY’ model, the values for GNa, GCa, GBK and GSK were directly extracted from Warashina’s model. To build the ‘SHR’ model, i) we assumed that the transcriptional changes equally mirror protein expression level and ii) because the electrical firing pattern results from the concomitant contribution of NaV, CaV and Ca^2+^-dependent K^+^ channels, we chose to change the conductance values globally, rather than individually (see data in Figure S7 for individual changes in GNa, GCa or GSK/BK). The conductance values implemented into the model were extrapolated from the changes in the expression level of the most abundant transcript for NaV, CaV and Ca^2+^-dependent K^+^ channels, that are NaV1.3 (1.4-fold), CaV1.3 (2-fold) and KCa1.1/KCa2.3 (1.7-fold) (Figure 6B). Depolarizing voltage steps (500 ms duration) of gradually increased stimulation intensities (10 to 35 μA/cm^2^) were injected into the ‘WKY’ and ‘SHR’ models. Representative APs evoked for three stimulation intensities are illustrated in Figure 7A. For low intensity stimulation (≤23 μA/cm^2^), the ‘SHR’ model generates more APs than the ‘WKY’ model, as observed in electrophysiological recordings (Figure 4A, upper charts). Consistent with the data obtained in the loose-patch configuration, high stimulation intensities (>23 μA/cm^2^) triggered less APs in the ‘SHR’ model. The AP frequency is maximal for stimulations >30 μA/cm^2^ and saturates at 24-26 Hz for ‘WKY’ and 12 Hz for ‘SHR’ models (Figure 7B). As such, and despite the fact that the ‘SHR’ model was compiled on the basis of transcriptional changes, our data suggest that a coordinate increase in NaV, CaV and Ca^2+^-activated K^+^ channels can account for the reduced excitability of SHR chromaffin cells, that occurs in response to robust depolarizations.

**Figure 7:**
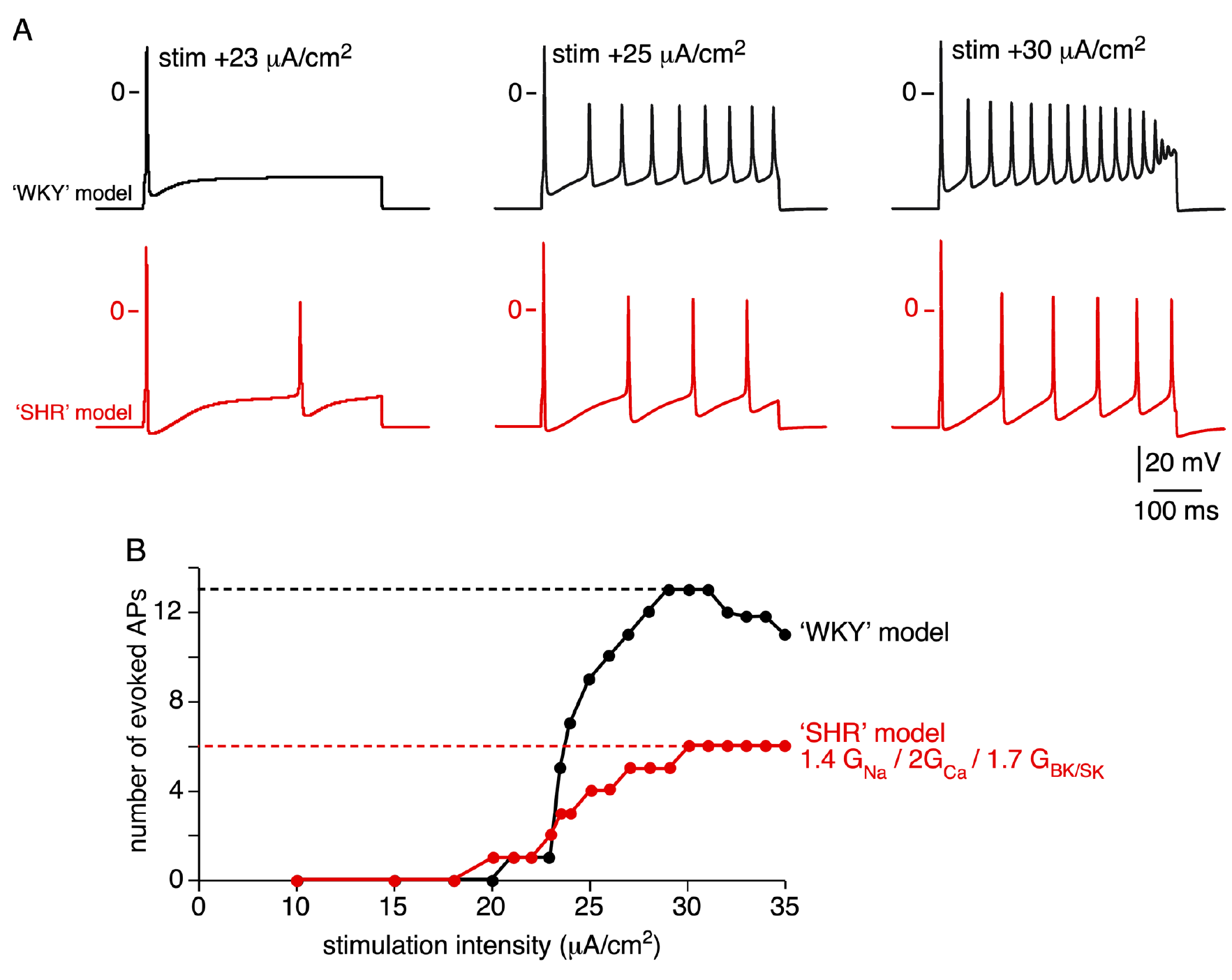
Numerical simulation of the electrical firing of WKY and SHR chromaffin cells: reduced excitability in SHR model in response to sustained depolarizations. ‘WKY’ and ‘SHR’ models were built from the rat chromaffin cell numerical simulation developed by Warashina and Ogura ^30^. The ‘SHR’ condition was simulated by added a multiplier (derived from the transcriptional changes) to Na^+^, Ca^2+^ and Ca^2+^-dependent K^+^ conductances, as such the ‘SHR’ model = 1.4 GNa, 2GCa and 1.7 GSK/BK ‘WKY’ model. The stimulation intensity was gradually increased from 10 to 35 μA/cm^2^ (500 ms depolarizing steps). **A.** Representative APs extracted from ‘WKY’ (black traces) and ‘SHR’ (red traces) models plotted for three stimulation intensities. **B.** Pooled data illustrating the reduced cell excitability in the ‘SHR’ model, in response to robust depolarizations (>23 μA/cm^2^).

### Changes in cholinergic synaptic transmission in hypertensive rats

Beside chromaffin cell excitability, cholinergic synaptic transmission at the splanchnic nerve-chromaffin cell junction is a key element of the stimulus-secretion coupling in the adrenal medulla. We therefore monitored the behaviour of cholinergic synapses by recording spontaneous excitatory post-synaptic currents (sEPSCs) in chromaffin cells voltage-clamped at-80 mV, as described ^31^. Consistent with previous studies ^32–34^ and reflecting the large number of nerve fibers cut during the slicing procedure ^35,36^, few chromaffin cells exhibited sEPSCs in standard 2.5 mM K^+^-containing saline (36.7%, n = 18/49 in WKY rats *versus* 23.1%, n = 12/52 in SHRs, p = 0.1909, Fisher’s exact test). Figure 8A illustrates representative charts of sEPSCs recorded in WKY (upper trace) and SHR (lower trace) chromaffin cells voltage-clamped at -80 mV, in response to a 80 mM KCl puff to increase the number of synaptic events. Mean sEPSC amplitude did not significantly differ between SHRs and WKY rats (133.1 ± 76.3 pA, n = 7 cells in SHRs and 106.3 ± 53.7 pA, n = 8 cells in WKY rats, p = 0.4634, Mann-Whitney test). Conversely, sEPSC frequency was significantly lower in SHR chromaffin cells (2.5 ± 2.2 Hz, n = 7 cells in SHRs *versus* 5.3 ± 2.2 Hz, n = 8 cells in WKY rats, p = 0.0093, Mann-Whitney test). sEPSCs kinetic parameters were also analyzed (Figure 8B). No difference was observed on the activation phase (rise time of 2.04 ± 0.25 ms, n = 8 cells and 2.18 ± 0.30 ms, n = 7 cells for WKY and SHR rats, respectively, p = 0.3357, Mann-Whitney test). By contrast, when the decay phase was fitted by a single exponential curve ^32,37^, a significant increase of the time constant was observed in SHRs (τ = 15.58 ± 2.19 ms, n = 8 cells for WKY rats *versus* 23.82 ± 5.25 ms, n = 7 cells for SHRs, p = 0.0037, Mann-Whitney test). The quantal analysis of sEPSCs did not show difference between SHRs and WKY rats (16.8 ± 4.4 pA, n = 8 cells for WKY rats and 20.5 ± 6.60 pA, n = 7 cells for SHRs, p = 0.2676, Mann-Whitney test, Figure 8C). Collectively, these data argue for synaptic activity changes occurring both at pre- and postsynaptic sites.

**Figure 8:**
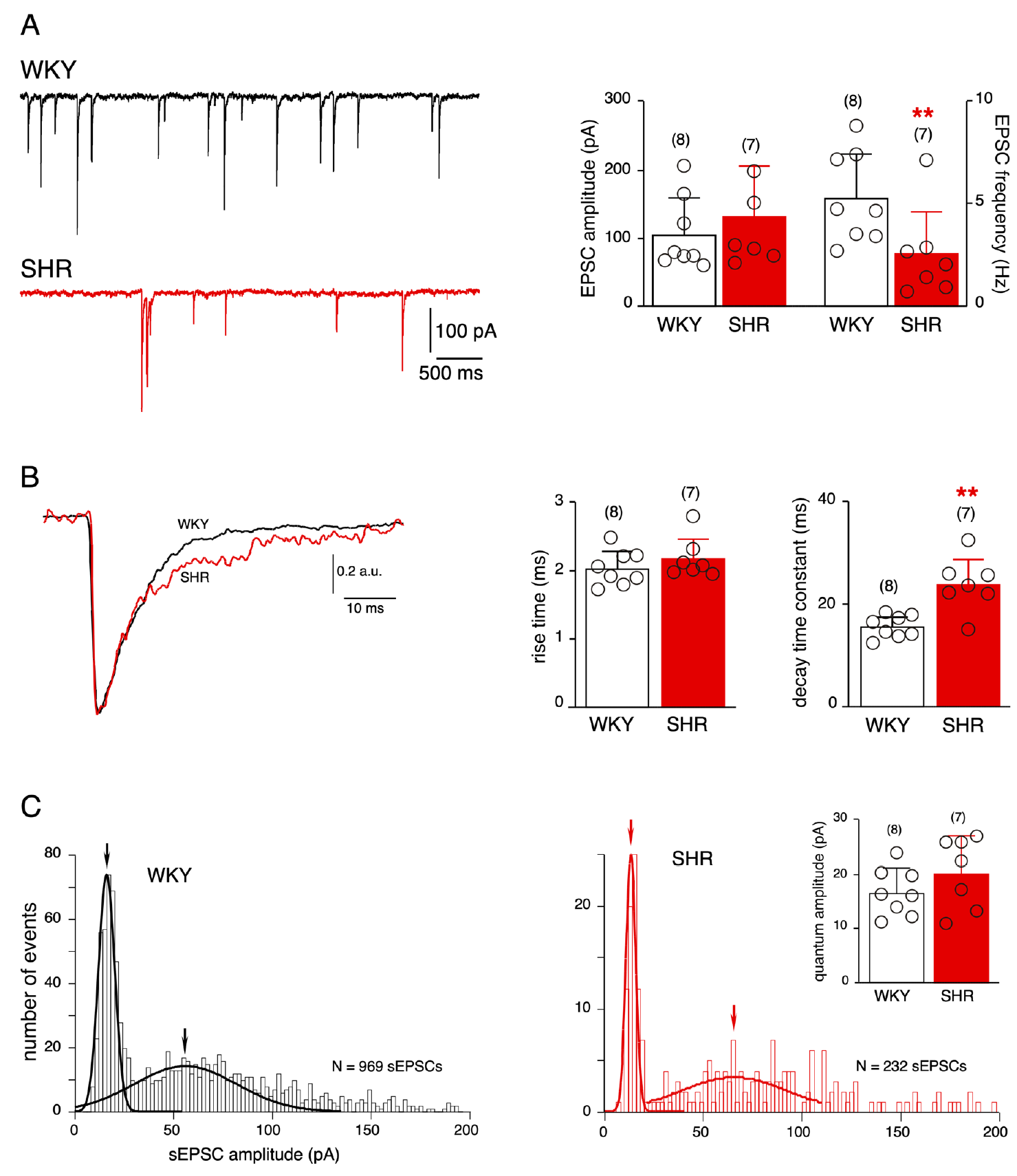
Changes in the excitatory cholinergic synaptic neurotransmission between splanchnic nerve endings and chromaffin cells in SHRs. **A.** Typical chart recordings of excitatory post-synaptic currents (EPSCs) recorded in WKY (upper trace) and SHR (lower trace) chromaffin cells voltage-clamped at -80 mV, in response to an 80 mM KCl puff. The analysis of synaptic transmission shows a significant decrease in EPSC frequency in SHRs (right histograms). **B.** In regard to EPSC kinetics, superimposed normalized WKY and SHR EPSCs and associated data histograms illustrate a significant elongated EPSC decay time in SHRs. **C.** Analysis of EPSC quantal size. Histograms (2 pA bin) illustrate of the distribution of sEPSC amplitudes in a WKY (left) and a SHR (right) chromaffin cells. Quantal size was estimated from the mean value of the first Gaussian curve fitted to the amplitude distribution histogram. As summarized from the 8 WKY cells and the 7 SHR cells in which the quantal analysis was performed, the EPSC quantal size does not differ between the two strains.

Excitatory postsynaptic events at the rat splanchnic nerve terminal-chromaffin cell junction result from the co-activation of several nicotinic acetylcholine receptor (nAChR) subtypes ^28,31,32,37^. A change in the expression and/or composition of nAChRs could account for the change in sEPSC decay time constant observed in SHRs. To address this issue, the expression level of transcripts encoding α3 (*Chrna3*), α4 (*Chrna4*), α5 (*Chrna5*), α7 (*Chrna7*),β2 (*Chrnb2*) and β4 (*Chrnb4*) subunits was quantified by real-time PCR (Figure 9). The expression of genes encoding α3 and α7, two subunits dominantly engaged in the synaptic transmission, did not change between SHRs and WKY rats. By contrast, the transcripts encoding the two auxiliary α4 and α5 subunits were differently expressed, with a significant increase for α4 (5.7-fold, p = 0.0043, Mann-Whitney test) and decrease for α5 (0.57-fold, p = 0.0043, Mann-Whitney test) in SHRs. For the β subunit family, the expression of β4, the main β subunit expressed in rat chromaffin cells significantly decreased in SHRs (0.48-fold, p = 0.0043, Mann-Whitney test). *Chrnb2* expression remained unchanged. Taken together, these findings indicate that nAChR subunits are transcriptionally remodeled in SHRs. Assuming subsequent modifications at the protein level, this plasticity may contribute to the modifications observed in sEPSC kinetics.

**Figure 9:**
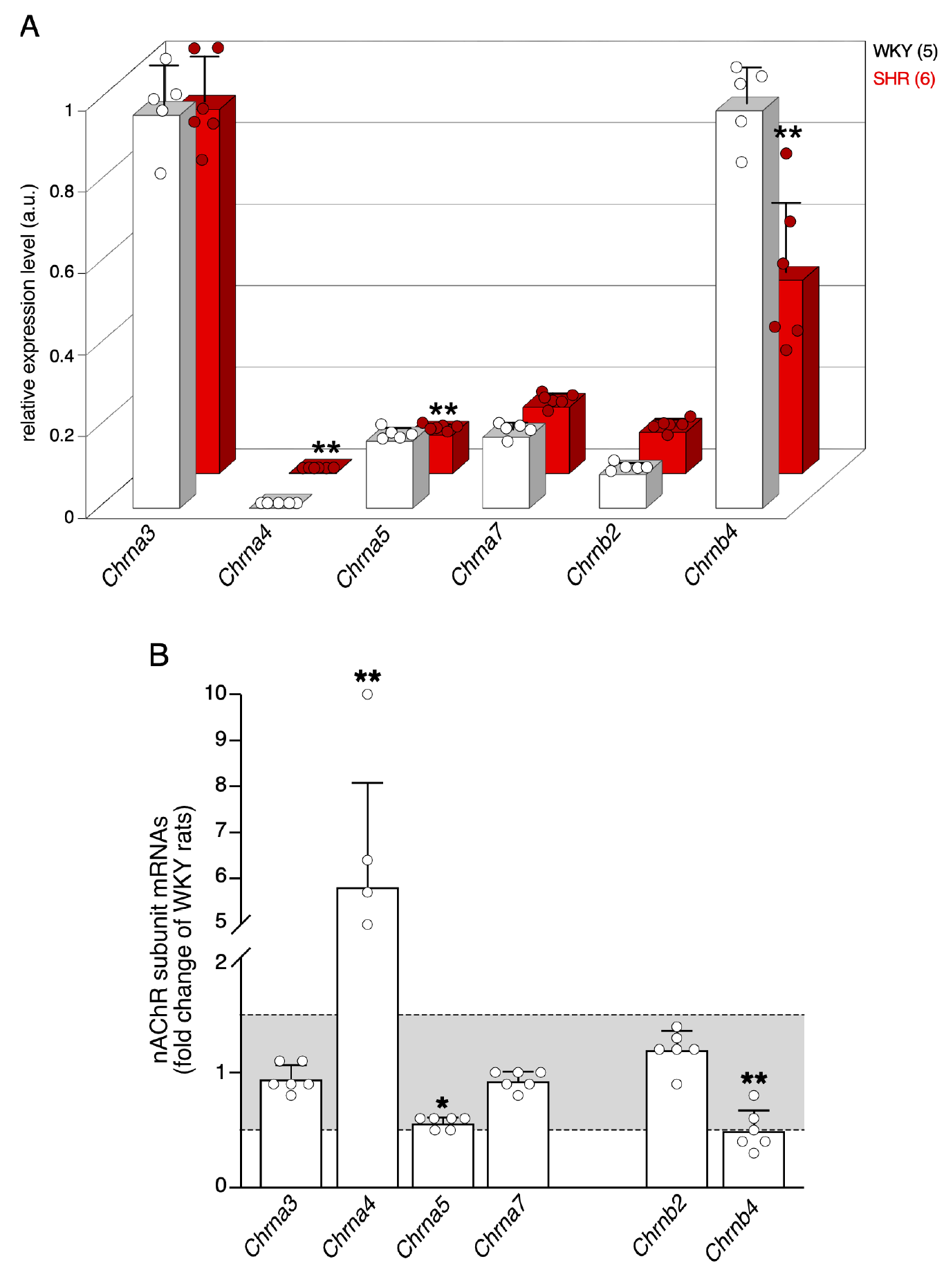
Remodeling of transcripts encoding nAChR subunits in SHRs. The changes in mRNA expression level of nAChR subunits were assessed by real-time RT-PCR, in macrodissected WKY (5) and SHR (6) adrenal medullary tissues. **A.** 3D-histograms illustrating the relative expression levels of four α (*Chrna3*−5 and *Chrna7,* encoding α3, α4, α5 and α7, respectively) and two β (*Chrnb2 and Chrnb4*, encoding β2 and β4, respectively) subunit genes, in the two rat strains. **B.** Fold changes in SHRs, as compared to WKY rat. Significant changes occur for *Chrna4* (5.8-fold), *Chrna5* (0.6-fold) and *Chrnb4* (0.5-fold). Fold change values were determined according to Livak’s method ^85^. The Shapiro-Wilk test was used to analyze the normality of data distribution, and parametric or non-parametric unpaired tests were used when appropriate. Fold changes between x0.5 and x1.5 (grey area) are considered irrelevant.

### Reduced gap junctional coupling between chromaffin cells in hypertensive rats

In the rat adrenal medullary tissue, gap junction-mediated intercellular communication between chromaffin cells is an additional pathway involved in the regulation of stimulus-secretion coupling ^15,17,19^. Because adrenomedullary gap junctional coupling can remodel in physiological/physiopathological conditions ^18,28,33,34,37–39^, we compared its status between SHRs and WKY rats. To investigate whether gap junctional coupling is modified in hypertensive rats, we imaged Lucifer yellow diffusion between chromaffin cells by confocal microscopy (Figure 10). As illustrated in panel 10A, the percentage of Lucifer yellow-coupled cells was lower in SHRs (18%, n = 7/38 cells in SHRs *versus* 33%, n = 11/33 cells in WKY rats, p = 0.1789, Fisher’s exact test). Consistently, the expression of connexin 43 (Cx43), the main connexin expressed in the rat adrenal medullary tissue ^15,38^, is significantly diminished in SHRs, both at the transcriptional (Figure 10B, left histogram) and protein (Figure 10C) levels. The expression of transcripts encoding Cx36 however did not change in hypertensive rats (Figure 10B, right histogram) These results, in addition with an increased input resistance (see Table S1), are consistent with a reduced gap junctional communication in SHRs. To go further into the mechanisms underlying the reduced expression of Cx43 in SHRs, we addressed the involvement of posttranslational changes such as stability of gap junctional plaques. Anchored gap junctions at the plasma membrane bind to scaffolding proteins, such as zonula occludens-1 (ZO-1). ZO-1 has been shown to interact with various connexins, including Cx43 ^40^. As shown by the densitometric analysis of immunoblots, ZO-1 expression remains unchanged in SHRs (Figure 10C), indicating that the impairment in Cx43 stability at the plasma membrane is unlikely the cellular mechanism whereby Cx43 expression is reduced in SHRs.

**Figure 10:**
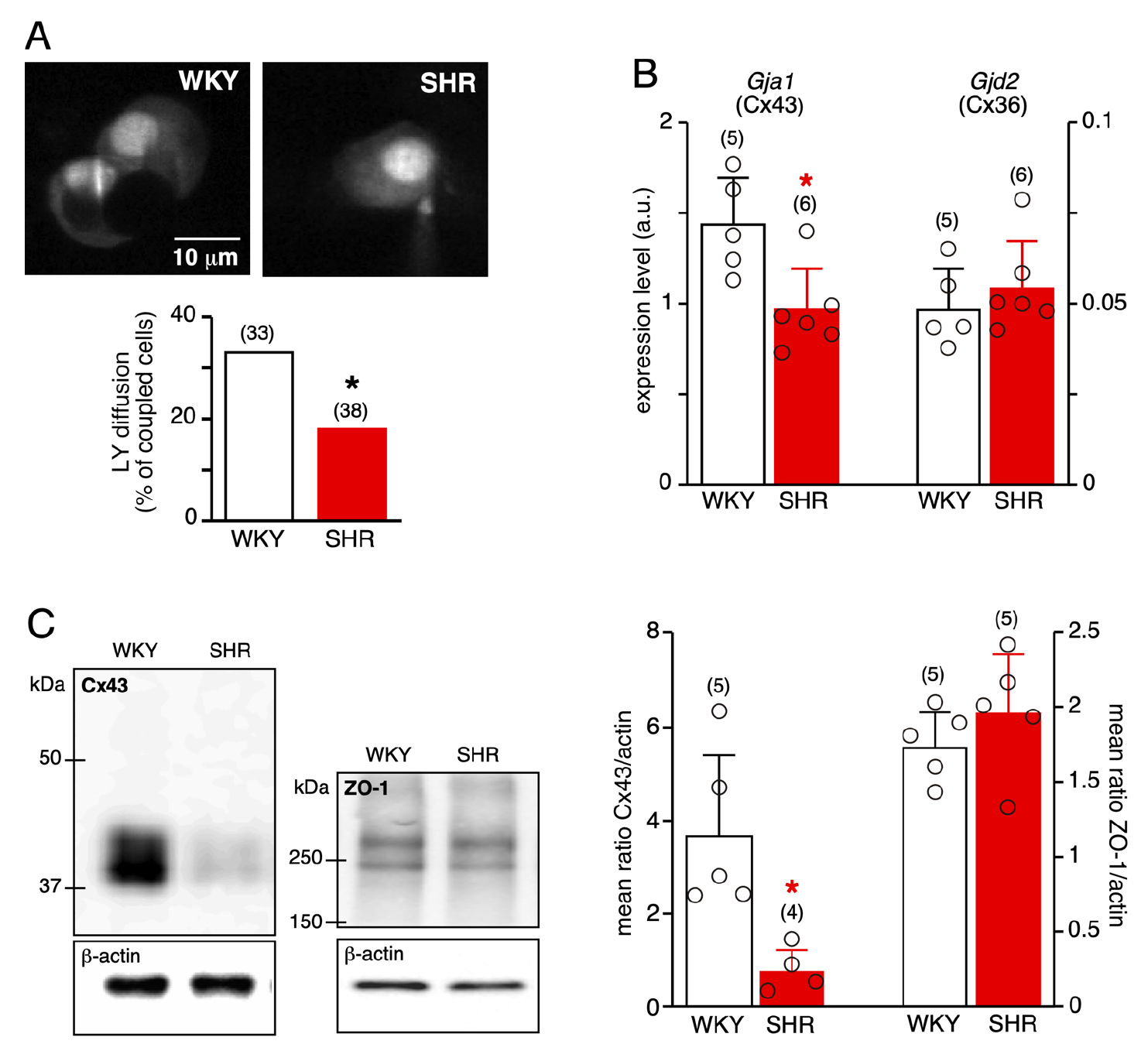
Attenuated gap junction-mediated intercellular communication in SHRs. **A.** Reduced Lucifer yellow (LY) diffusion between SHR chromaffin cells. LY was introduced into chromaffin cells using patch pipettes. Dye diffusion was imaged 15 minutes after patch disruption. Less than 20% of SHR chromaffin cells are dye-coupled, as compared to more than 30% in WKY rats. **B.** Decreased expression level of *Gja1* (encoding Cx43), but not *Gjd2* (encoding Cx36) in SHRs, assessed by real-time RT-PCR from macrodissected medullary tissues. **C.** Western blots and associated pooled data histograms of Cx43 and ZO-1, an associated protein. Cx43 expression, but not ZO-1, is significantly down-regulated in SHRs.

## Discussion

This study describes the functional remodeling operated by adrenal chromaffin cells in response to chronic elevated blood pressure. All observed changes converge toward a less efficient stimulus-secretion coupling, especially in response to robust stimulations. gis may reflect a shielding mechanism dedicated to keep the hypertensive organism safe of the deleterious effects on huge CA secretion episodes.

In addition to the functional changes unveiled between ’normotensive’ and ’hypertensive’ chromaffin cells, this study also extends the basic knowledge on rat chromaffin cell excitability *in situ*. The analysis of AP firing revealed the existence of two spiking patterns (regular and bursting), as previously reported in mice ^18,23,41^. As in the mouse, regular and bursting discharges coexist in a same cell. Because APs generated in bursts are more efficient than AP trains fired at constant frequency to evoke CA secretion ^26,42^, identifying the cellular mechanism(s) promoting the switch between the two spiking patterns would be therefore of great interest. In this context, ion channels involved in setting up the resting membrane potential appear as interesting targets to investigate ^23,43^. In addition, although we found no sharp difference in the occurrence of regular and bursting patterns between WKY rats and SHRs, further experiments are required to fully address this issue, especially in response to adrenomedullary secretagogues (neurotransmitters, neuropeptides…).

### Decreased chromaffin cell excitability in response to robust depolarization/stimulation

The most functionally relevant feature of SHR chromaffin cells is their reduced AP firing in response to sustained stimulations. gis likely results from a coordinated upregulation of Na^+^, Ca^2+^ and Ca^2+-^dependent K^+^ channels, as observed both experimentally with the enhanced expression level of their respective transcripts and *in silico* in a chromaffin cell model. At the single AP level, AP waveform in SHR chromaffin cells is modified as such the amplitude is decreased and the half-width is increased. Because the AP waveform results from the concomitant contribution of many ion channels ^44^, discussing each channel separately would give biased conclusions. We therefore focused the discussion on the most striking effects that is the AP waveform change and reduced ability of SHR chromaffin cells to trigger sustained firing during robust depolarizations. In chromaffin cells, as in most neurons, NaV channels critically shape the AP waveform ^45^. gis is of particular importance because changes, even subtle, in spike shape can drastically affect Ca^2+^ channel recruitment and therefore ensuing Ca^2+^transients, and finally CA secretion. Interestingly, the more remarkable change in ion channel expression in SHRs targets NaV1.7, a sodium channel involved in regulating chromaffin secretory function ^46^. As reported ^26^, the reduced NaV channel availability (through NaV1.3 and NaV1.7) switches cell firing from a repetitive AP firing mode to a bursting activity, boosting thus hormone exocytosis. At rest, NaV channels tonically inhibit burst firing and would contribute to maintain a low basal CA secretion. In SHRs, the significant increase of NaV1.7 would help retaining chromaffin cell to spontaneously fire in a tonic mode, preserving the gland of huge CA secretion. gis is consistent with our data showing that basal NE and E secretion is not enhanced in SHRs (even decreased for NE). The concomitant increased expression of NaV1.7, CaV1.3 and KCa channels in SHR chromaffin cells is consonant with data reporting that i) L-type Ca^2+^ channel activity can upregulate NaV channels in endocrine cells ^47,48^ and ii) CaV1.3 and BK/SK channels are functionally coupled in chromaffin cells ^41,45,49^. Long-term changes in NaV channel expression have been also described in endocrine tissues in response to stressful situations ^50^. Collectively, these findings prompt us to propose the provisional sequence of events, that is an enhancement of L-type CaV channels and ensuing cytosolic calcium followed by a subsequent enhancement of NaV and KCa channels as a shielding mechanism to brake CA secretion, especially in response to prolonged or repetitive stimulations. We have focused here on voltage-gated ion channels involved in AP firing, but other ion channels, K^+^/Na^+^ background/leak channels operating at resting membrane potential in particular, are also qualified to modify cell excitability ^23,43,51^ and would therefore be interesting to investigate.

### Weakened synaptic transmission in SHRs

In SHRs, the excitatory neurotransmission between splanchnic nerve and chromaffin cells is pre- and postsynaptically remodeled. According to the quantal theory of neurotransmitter release ^52^, the reduced number of sEPSCs in response to a sustained stimulation may relate to failures eventuating on presynaptic components. Additionally, and supported by changes occurring postsynaptically at nAChRs, the incoming signals would be less efficiently integrated by hypertensive cells. In chromaffin cells, the prevalent nAChR compositions encompass α3β4 and α3β2 subunits with the addition of α5 ^53^. Because the β4 subunit modulates the density α3β4 receptors at the plasma membrane ^54^, the decreased expression level of β4 mRNA in SHRs may reduce the membrane expression of α3β4 nAChRs and attenuate the neurotransmission at splanchnic nerve-chromaffin cell synapses. When incorporated in a nAChR, α5 subunit alters biophysical and biochemical properties of the channel ^55,56^. Of a particular interest for a neurosecretory cell is the less robust Ca^2+^ permeability of α3 nAChRs in the absence of α5 ^56^. If we assume that the reduced expression level of α5 transcripts in SHRs could favor α3β2/ α3β4 combinations to the detriment of α3β2α5/α3β4α5, it is likely that the resulting Ca^2+^ entry through activated nAChRs could be less robust, and could contribute to reduce hormone secretion. Both α4β2 and α4β4 combinations build functional nAChRs, although the current properties differ between the two combinations ^57^. The down-regulation of β4 in SHRs suggests that the prevalent combination in hypertensive rats could be α4β2 rather than α4β4. Assuming this, the current flowing through α4β2 nAChRs would then display a lower amplitude and a faster desensitization ^57^. Collectively, the transcriptional remodeling of nAChR subunit expression observed in SHRs converges towards a less efficient cholinergic coupling between nerve endings and chromaffin cells in hypertensive animals.

### Reduced gap junctional coupling in SHRs

Gap junctional signaling between chromaffin cells is a part of regulatory mechanisms involved in stimulus-secretion coupling ^16,17,19,20^. Although we did not directly assess the junctional current, the reduced LY diffusion and the decreased Cx43 protein expression level strongly suggest an attenuated gap junctional communication in SHRs. Changes in chromaffin cell input resistance and capacitance, two parameters impacting the transmission strength and therefore the propagated signals ^58^, also support this statement. Given that gap junction coupling contributes to CA secretion ^15,18,20,38^, a decreased intercellular communication could be a suitable mechanism whereby the adrenal medulla would damper its secretory process, especially in response to harmful challenges (*i.e.* repetitive stimulating episodes).

### Reduced CA release in SHRs in response to sustained cholinergic stimulations

Our results argue for a decrease in the stimulus-secretion coupling efficiency in SHRs. E and NE secretion is attenuated in SHRs, especially in response to high ACh concentrations. At rest and for lower ACh concentrations, only adrenal NE outflow is decreased, as previously reported^59^. gis suggests that the ‘hypertensive’ environment (circulating levels of CA and other adrenomedullary hormones/factors, adrenocortical secretions…) differentially impacts the mechanisms involved in regulating NE and E release. However, besides their difference in secretory competence, the hypertensive and normotensive medullary tissues are equally readily responsive to a cholinergic stimulus, as shown by equivalent stimulation ratios. Interestingly, they differ in the E:NE ratios in response to a cholinergic stimulus. Whereas in WKY rats, E:NE ratio increases gradually with ACh concentration (due to enhanced E release, as expected in a healthy adrenal medulla), it remains constant in SHRs, indicating a dysregulation in the secretory process of E There are many targets (secretory granules, exocytosis…), and further experiments are needed to identify the impaired processes.

Several explanations can account for the decreased stimulus-secretion coupling efficiency in SHRs. First and supported by our electrophysiological data, SHR chromaffin cell less robustly fire upon strong depolarizations. gis, in association with the remodeling of nAChR subunits, likely contributes to lessen electrical activity- and cholinergic-evoked [Ca^2+^]i rises, and subsequent CA exocytosis. Second, downstream changes may also occur, targeting for example CA biosynthesis and/or intra-adrenal content. Regarding this, the literature is puzzling, that is either an increase or a decrease in activity/mRNA expression level of TH, DβH and PNMT in adult SHRs have been described ^60–65^. Similarly, the intra-adrenal CA content in adult SHRs has been reported to be either decreased, increased or unchanged ^62,65–67^. Along the same line, the secretory granules could also be impacted. Of a particular interest are the granins, major components of chromaffin granules critically involved in granulogenesis, granule cargo and exocytosis ^68,69^. By regulating circulating CA levels, granins and their derived peptides are potent modulators of blood pressure ^70–72^. In adult SHRs, the increased adrenal content of chromogranin A ^62,67^ is likely one of the mechanisms contributing to elevated blood pressure.

### Physiological and/or pathological consequences of attenuated stimulus-secretion coupling in adult SHRs: a shielding mechanism of the adrenal medullary function and/or impairment of the stress response?

The question is open to debate. From a physiological point of view, our finding of a reduced stimulus-secretion coupling in the SHR adrenal medulla is functionally relevant and helps to decode how chromaffin cells behave to cope with a hypertensive environment, and more generally with a chronic hyperactivity as observed during a sustained splanchnic nerve firing activity evoked by prolonged stress episodes. The reduced CA release would also preserve from exhaustion of the secretory process, maintaining therefore the tissue still competent for further stimulations. In the same context, given the importance of CA in the control of hemodynamic properties (heart rate, cardiac output, blood pressure, vascular tone), the reduced NE outflow observed at rest and for lower ACh concentrations could be physiologically relevant to mitigate the deleterious consequences of a hypertensive environment. Indeed, by activating α-adrenoceptors, NE increases peripheral arterial resistance and therefore blood pressure. E by acting on both α- and β-adrenoceptors has also a hypertensive effect, but in a lesser extent compared to NE. Beyond their hemodynamic properties, plasma CA, when chronically elevated, can promote pathophysiological conditions including inflammation, metabolic disorders, and organ failures ^73^. In this context, any mechanism aimed at reducing plasma CA levels can be considered protective.

In the same line, we can also propose that the remodeling of the adrenal medulla in hypertensive animals could prevent excessive energy expenditure at both tissue and organism levels. CA synthesis and secretion are energy-consuming processes ^74^, so we speculate that weakening the competence of the adrenal medulla to release catecholamines could limit this energy cost. In addition, since elevated catecholamine secretion is known to induce an increase of energy expenditure mediated by metabolism stimulation ^75^, the remodeling we describe here would be beneficial to limit catecholamine secretion.

Adrenal medulla-driven adaptive mechanisms are decisive for the ability of an organism to cope with stress (^76^ for a recent review) and any dysfunction or imbalance in one of these mechanisms can jeopardise its survival. One issue, not investigated here, would be to evaluate whether and how SHRs and more generally hypertensive animals adapt to stress (acute/chronic). Does the ’hypertensive’ adrenomedullary tissue undergo remodeling of chromaffin cell excitability, synaptic cholinergic/peptidergic neurotransmission and gap junctional coupling? In this context, PACAP, a major stress transmitter at the splanchnic nerve-chromaffin cell synapse ^25^, is certainly an avenue to explore. Our data showing an alteration of the electrical response to PACAP together with i) the previously reported effect of PACAP on *Pnmt* gene expression ^77^ and ii) an increased expression of *Pnmt* mRNA and protein levels in SHRs ^64,78^ suggest altered (defective?) adrenergic transduction pathways/function in SHRs.

As such, the adrenal stimulus-secretion coupling is a dynamic process, continuously remodeled according to physiological or pathophysiological conditions^28,79^. Whereas acute CA needs (as required during stressful situations) are associated with an increased stimulus-secretion coupling efficiency ^18,34,38^, a long-lasting impregnation of the adrenal medullary tissue with high circulating CA levels (as observed during hypertension development for example) would correlate with a reduced stimulus-secretion coupling competence.

## Materials and Methods

### Ethical approval and animals

Animals were housed in groups of 3-4 par cages (standard sizes according to the European animal welfare guidelines 2010/63/EU) and maintained in a 12h light/dark cycle, in stable conditions of temperature (22°C) and humidity (60%). Food and water were provided *ad libitum*. All procedures in this study also conformed to the animal welfare guidelines of the European Community and were approved by the French Agriculture and Forestry Ministry (authorization numbers/licences 49-2011-18, 49-247, A49007002 and D44015) and by the regional ethic committee (authorization CEEA.2011.12 and APAFIS#2017072117413637).

Data were collected from 16- to 20-week-old spontaneously hypertensive male rats (SHRs) and from their control age-matched normotensive male Wistar-Kyoto (WKY) rats (SHR/KyoRj and WKY/KyoRj, Janvier Labs, Le Genest-St-Isle, France). The body weight did not significantly differ between SHRs (401.4 ± 33.3 g, n = 38) and WKY rats (417.2 ± 43.6 g, n = 47, p = 0.0689, unpaired t test, Figure S1A). As a consequence of the arterial hypertension, the heart weight was statistically higher in SHRs (1.51 ± 0.23 g, n = 38 *versus* 1.22 ± 0.10 g, n = 47 in WKY rats, p<0.0001, unpaired t test, Figure S1B). Accordingly, the heart/body weight ratio (x100) was also significantly higher in SHRs (0.38 ± 0.06, n = 38 *versus* 0.29 ± 0.02, n = 47 in WKY rats, p<0.0001, unpaired t test, Figure S1C).

### Adrenal slice preparation

Acute slices from SHRs and WKY rats were prepared as described ^15,80^. Briefly, after removal, the adrenal glands were kept in ice-cold saline for 2 min. Before slicing, a gland was desheathed of the surrounding fat tissue and was next glued onto an agarose cube and transferred to the stage of a vibratome (HM 650V vibrating blade microtome, Microm Microtech, Francheville, France). Slices (200 μm thickness for the electrophysiological recordings and 150 μm thickness for CA secretion assay) were cut with a razor blade and transferred to a storage chamber maintained at 37°C, containing Ringer’s saline (in mM): 130 NaCl, 2.5 KCl, 2 CaCl2, 1 MgCl2, 1.25 NaH2PO4, 26 NaHCO3, 12 glucose and buffered to pH 7.4. The saline was continuously bubbled with carbogen (95% O2/5% CO2).

### Electrophysiology

All electrophysiological recordings were performed in acute slices. Adrenal slices were transferred to a recording chamber attached to the stage of a real-time confocal laser scanning microscope (LSM 5Live, Zeiss) equipped with an upright microscope (Axio Examiner, Zeiss, Le Pecq, France) and continuously superfused with Ringer’s saline at 34°C. All experiments were performed using the patch-clamp technique and electrophysiological signals were acquired using an EPC-10 USB Quadro patch-clamp amplifier (HEKA Electronik, Lambrecht/Pfalz, Germany) and PATCHMASTER software. Signals were sampled at 10 Hz and analyzed with FITMASTER (HEKA Electronik, Germany). For whole-cell recordings, pipettes were pulled from borosilicate glass and filled with the following internal solution (in mM): 140 potassium (K)-gluconate, 5 KCl, 2 MgCl2, 1.1 EGTA, 5 Hepes, 4 MgATP, 0.3 NaGTP and titrated to pH 7.2 with KOH. Osmolarity was adjusted at 295 mOsm with K-gluconate and pipette resistance was 5-8MΩ. Voltage was corrected for the junction potential of about 13 mV. Pipette and cell capacitances were fully compensated and the series resistance was compensated at 75-80%. Membrane potential was recorded in the current-clamp mode and filtered at 3 kHz. The AP properties were examined by injecting current pulses of 500 ms duration from -50 to +60 pA at 1 Hz. The evoked AP analysis was done using Mini Analysis software (Synaptosoft Inc. Fort Lee, NJ, USA). For extracellular recordings of spontaneous AP currents in the loose-patch configuration, pipettes were pulled to a resistance of 5-10 MΩ when filled with the following saline (in mM): 130 NaCl, 2.5 KCl, 2 CaCl2, 1 MgCl2, 10 HEPES, 10 glucose and buffered to pH 7.4 with NaOH. Osmolarity was adjusted at 295 mOsm with NaCl. The liquid junction potential was approximately 0 mV. Once the tip of the pipette was positioned at the surface of a chromaffin cell, a minimal suction pressure was applied (seal resistance <500 MΩ) and the electrical activity was recorded in the voltage-clamp mode (0 mV) of the loose cell-attached configuration ^81^. gis method allows investigation of membrane excitability under physiological conditions and stable recordings of firing rate can therefore be obtained ^82,83^. Regarding the extracellular experiments with PACAP, after a 2-5min stabilisation period, we initiated recordings for 5 minutes (control condition). Using a Pressurized Drug Applicator (NPI electronic, Bauhofring, Germany) attached to a glass pipette, PACAP was then puffed near the recorded cell for 5 minutes (PACAP condition). The results were represented as a percentage of the AP frequency following PACAP application relative to the control condition. gree different doses of PACAP were used (0.1, 1 and 10 µM) on distinct slices and chromaffin cells. Spontaneous excitatory postsynaptic currents (sEPSCs) were recorded in chromaffin cells voltage-clamped at -80 mV and were filtered at 1 kHz, as previously described ^31^. To increase the percentage of chromaffin cells exhibiting sEPSCs, recordings were performed by applying a 80 mM KCl puff at the vicinity of the recording cells ^32,35^. In some cells, a quantal analysis of sEPSCs was performed. Only single events were selected for construction of amplitude histograms. The histograms were inspected for the presence of peaks and a corresponding number of Gaussians was then fitted by nonlinear regression using GraphPad Prism 5.0 software (San Diego, USA). Quantal size was estimated from the mean value of the first Gaussian curve fitted to the amplitude distribution histogram. Normalized sEPSC decays were fitted by a single exponential using the Simplex fit algorithm based on the sum of squared errors SSE. Only fits with a SSE<0.001 were taken into consideration. The analysis was done using Mini Analysis software.

### Mathematical model of chromaffin cell electrical firing

Chromaffin cell electrical activity was modeled using the chromaffin cell simulation developed by Warashina and Ogura ^30^. The numerical model was adapted to the JSim software ^84^, available at https://www.imagwiki.nibib.nih.gov/physiome/jsim. The time adaptive integration using the Radau method (dt = 0.01 ms) was used to give stable solutions to ordinary differential equations. Conductance changes between SHRs and WKY rats were extrapolated from the changes observed in the expression level of the most abundant transcript for Na^+^, Ca^2+^ and Ca^2+^-dependent K^+^ channels, that are Nav1.3 (1.4-fold change), Cav1.3 (2-fold change) and KCa1.1/KCa2.3 (1.7-fold change).

### Dye transfer assay

The fluorescent dye LY (Lucifer yellow-CH, dilithium salt, 1 mM in internal solution) was introduced into chromaffin cells using patch pipettes. The probability of LY diffusion was expressed as a ratio corresponding to the number of injected cells that show dye transfer to adjacent cells over the total number of injected cells, as previously described ^38^. Cells were viewed with a 63x/1.0 NA Plan-Apochromat water immersion objective (Zeiss). Dye transfer between gap junction-coupled cells was imaged with a real-time confocal laser-scanning microscope (LSM 5Live, Zeiss) equipped with a diode laser 488 nm (100 mW). The extent of LY diffusion was estimated by counting the number of neighboring cells that received dye in 15 min.

### Quantification of mRNA expression levels by real-time PCR

Total RNA was extracted from macrodissected adrenal medulla using the RNeasy^®^ Micro extraction kit (Qiagen, Courtaboeuf, France), as previously described ^18^. RNA (500 ng) was first reverse transcribed using the QuantiTect® Reverse Transcription kit (Qiagen) in a final volume of 10 μl. Real-time PCR analyses of the target genes and the reference genes *Gusb*, *Hprt* and *Gapdh* were performed using Sybr^®^ Green PCR master mix (Applied Biosystems, Foster City, CA) with 1:100 of the reverse-transcription reaction, and were carried out on an ABI 7500 SDS Real-Time PCR system (Applied Biosystems). After an initial denaturation step for 10 min at 95°C, the thermal cycling conditions were 40 cycles at 95°C for 15 s and 60°C for 1 min. Specificity of amplification was checked by melting curve analysis. Each sample value was determined from duplicate measurements. Expression of the target transcripts was normalized to the mean of the expression level of the three reference genes according to the formula E = 2^-(Ctmean[Target]-Ctmean[Reference])^, where Ctmean is the mean threshold cycle. The fold change values were determined according to Livak’s method ^85^. Primer sequences of target genes are given in Table S3 and the concentration used was 300 nM for all genes.

### Determination of Cx43 and ZO-1 protein expression

Cx43 and ZO-1 expression levels were achieved on macrodissected adrenal medulla (5 SHRs and 5 WKY rats). After decapsulation, the medullary tissue was separated from the cortex, quickly frozen and pulverized in liquid nitrogen. For each animal the right and the left medulla were pooled in a same sample. The samples were resuspended in lysis buffer (10 mM Tris-HCl, pH 7.4, 1 mM sodium orthovanadate, 10 mM NaF as phosphatase inhibitors and 1% SDS), supplemented with Mini complete protease inhibitors (Roche Applied Science, Laval, Quebec). The extracts were incubated 30 min at 4°C, sonicated and centrifuged for 20 min at 12000 g. The supernatant was then collected, and the protein concentration was determined using the Pierce BCA Protein Assays Kit (germo Fisher Scientific, Rockford, IL) Protein samples (25 μg) were separated by 10% SDS-PAGE (4-15% gradient for ZO-1 detection). Electrophoresed proteins were transferred onto a nitrocellulose membrane. Blots were then blocked with 5% milk in TBS (pH 7.4, 0.1% Tween 20) for 1 h at room temperature with gentle agitation. Blots were next incubated with polyclonal antibodies, a rabbit anti-Cx43 (1:500, germo Fisher Scientific) or a rabbit anti-ZO-1 (1:250, germo Fisher Scientific) in TBS-Tween 0.1% containing 5% milk, at 4°C overnight. Following washout, blots were incubated with secondary antibodies peroxydase conjugated for 1 h at room temperature. Proteins were visualized with a LAS-3000 imager (Fuji) using an ECL-Plus Chemiluminescence kit (germo Fisher Scientific). To ensure equal loading of protein samples, blots were stripped of connexin or ZO-1 antibodies and reprobed with an actin-specific monoclonal antibody (Ω-actin, 1:5000, Sigma). Intensities of Cx43 and ZO-1 bands were normalized to those of actin and quantified using the Las-3000 software.

### CA assays from adrenal slice supernatants

Chromaffin cell ability to secrete CA was assessed by monitoring E and NE released in slice supernatant, as previouly described ^22^. After a 5-min basal (B) condition, slices were challenged (S) with either ACh-containing saline or Ringer saline, during 5 min. For each slice, results were expressed as the stimulation ratio S/B. To calculate basal or stimulated amounts of secreted E and NE, the medulla surface was estimated for each slice. To achieve this, at the end of experiments, slices were bathed for 2 min in hematoxyline (1 g/l). The differential staining intensity between cortex and medulla allowed estimating the medulla surface, on the two slice faces. The volume was then subsequently calculated. After collection, samples were kept at -20°C during 2-3 days before use. E and NE were assayed by High Performance Liquid Chromatography (HPLC). A benzylamine pre-column derivatization was used to generate fluorescent benzoxazole derivatives ^86^, which were next separated by C18 reversed-phase HPLC (Vydac 218TP54 column, 5 mm, 4.6 mm i.d. x 250 mm; Waters Separations Module 2695; multiwavelength fluorescence detector Waters 2475) ^87^. The derivatization reactions were carried out using 20 μl of a standard solution of CA (L-epinephrine- and L-norepinephrine-L-bitartrate salts) or supernatant samples or water (blank) mixed to benzylamine (50 mM) and 3-cyclohexylaminopropanesulfonic acid (3.3 mM, pH 11) and potassium hexacyanoferrate III (1.7 mM). The resulting mixtures were incubated at 50°C for 20 min and 20 μl were loaded onto the reverse phase C18-column. Isocratic elution was performed with a mobile phase consisting of 10 mM acetate buffer (pH 5.5) and acetonitrile (65/35, v/v) run at 1 ml/min. The column temperature was maintained at 23°C. The detection was monitored at an excitation wavelength of 345 nm and an emission wavelength of 480 nm. To determine E and NE concentrations, the areas under the peaks of samples were compared to the peaks of the standard range used as external calibrator.

### Solutions and chemicals

Benzylamine, 3-cyclohexylaminopropanesulfonic acid, potassium hexacyanoferrate III, L-epinephrine-L-bitartrate salt, L-norepinephrine-L-bitartrate salt, acetonitrile, Lucifer yellow (dilithium salt), pituitary adenylate cyclase activating polypeptide-38 (PACAP-38) and acetylcholine chloride were purchased from Sigma-Aldrich (Saint-Quentin Fallavier, France).

### Statistical analysis

Statistics were performed with Prism 9 (version 9.4.1, GraphPad, San Diego, CA). Numerical data are expressed as the mean ± standard deviation. Differences between groups were assessed by using the non-parametric Mann-Whitney test. Unpaired Student’s t-test was used to compare means when appropriate. The non-parametrical Wilcoxon matched-pairs signed-rank test was used to compare two related samples or to compare a single sample to a theoretical value (1 or 100%). For comparisons of more than two groups, the non-parametric Kruskal-Wallis test was used. The Spearman’s rank correlation coefficient ρ was used to measure the relationship between paired data. Percentages were compared using a contingency table and the chi-square test or the Fisher’s exact test when appropriate. Differences with p<0.05 were considered significant, with *, p<0.05, **, p<0.01, ***, p<0.001 and ****, p<0.0001.

## Acknowledgments

We thank Drs. Michel G. Desarménien and Philippe Lory for helpful discussion in preparing the manuscript, Drs. Claire Legendre and Hélène Tricoire-Leignel for their technical advices in the molecular techniques, and Prof. Pascal Richomme for HPLC facility in his laboratory (Univ. Angers, SONAS, SFR QUASAV, Angers, France). gis work was supported by grants from Centre National de la Recherche Scientifique, Institut National de la Santé et de la Recherche Médicale, Fondation pour la Recherche Médicale, Région Pays de la Loire, Conseil Général de Maine et Loire and Angers Loire Métropole.

## Author contributions

V.P., J.P, B.T., J.B., F.D.N., P.F., C.G.-L., C.L. and N.C.G. conducted the experiments. D.B. and D.G. assisted with HPLC equipment.

V.P., J.P, B.T., J.B., F.D.N., C.G.-L., C.L. and N.C.G analyzed the data. N.C.G and C.L. designed the experiments. N.C.G. conceived the study and wrote the paper with input from D.H. and C.L.

## References

1. Tank, A.W., and Lee Wong, D. (2015). Peripheral and central effects of circulating catecholamines. Compr Physiol 5, 1–15. 10.1002/cphy.c140007.

2. Johnson, M.D., Grignolo, A., Kuhn, C.M., and Schanberg, S.M. (1983). Hypertension and cardiovascular hypertrophy during chronic catecholamine infusion in rats. Life Sci 33, 169–180. 10.1016/0024-3205(83)90410-1.

3. Schwartz, D.D., and Eikenburg, D.C. (1986). Cardiovascular responsiveness to sympathetic activation after chronic epinephrine administration. J Pharmacol Exp ger 238, 148–154.

4. Fregly, M.J., Kikta, D.C., greatte, R.M., Torres, J.L., and Barney, C.C. (1989), Development of hypertension in rats during chronic exposure to cold. J Appl Physiol 66, 741–749. 10.1152/jappl.1989.66.2.741.

5. Anderson, E.A., Sinkey, C.A., Lawton, W.J., and Mark, A.L. (1989). Elevated sympathetic nerve activity in borderline hypertensive humans. Evidence from direct intraneural recordings. Hypertension 14, 177–183. 10.1161/01.hyp.14.2.177.

6. Esler, M., Ferrier, C., Lambert, G., Eisenhofer, G., Cox, H., and Jennings, G. (1991). Biochemical evidence of sympathetic hyperactivity in human hypertension. Hypertension 17, III29–35. 10.1161/01.hyp.17.4_suppl.iii29.

7. Papanek, P.E., Wood, C.E., and Fregly, M.J. (1991). Role of the sympathetic nervous system in cold-induced hypertension in rats. J Appl Physiol (1985) *71*, 300–306. 10.1152/jappl.1991.71.1.300

8. Lim, D.Y., Jang, S.J., and Park, D.G. (2002). Comparison of catecholamine release in the isolated adrenal glands of SHR and WKY rats. Auton Autacoid Pharmacol 22, 225–232. 10.1046/j.1474-8673.2002.00264.x.

9. Friese, R.S., Mahboubi, P., Mahapatra, N.R., Mahata, S.K., Schork, N.J., Schmid-Schonbein, G.W., and O’Connor, D.T. (2005). Common genetic mechanisms of blood pressure elevation in two independent rodent models of human essential hypertension. Am J Hypertens 18, 633–652. 10.1016/j.amjhyper.2004.11.037.

10. Mathar, I., Vennekens, R., Meissner, M., Kees, F., Van der Mieren, G., Camacho Londono, J.E., Uhl, S., Voets, T., Hummel, B., van den Bergh, A., et al. (2010). Increased catecholamine secretion contributes to hypertension in TRPM4-deficient mice. J Clin Invest 120, 3267–3279. 10.1172/JCI41348.

11. Pak, C.H. (1981). Plasma adrenaline and noradrenaline concentrations of the spontaneously hypertensive rat. Jpn Heart J 22, 987–995. 10.1536/ihj.22.987.

12. Guerineau, N.C., Campos, P., Le Tissier, P.R., Hodson, D.J., and Mollard, P. (2022). Cell Networks in Endocrine/Neuroendocrine Gland Function. Compr Physiol 12, 3371–3415. 10.1002/cphy.c210031.

13. Douglas, W.W. (1968). Stimulus-secretion coupling: the concept and clues from chromaffin and other cells. Br J Pharmacol 34, 451–474. 10.1111/j.1476-5381.1968.tb08474.x.

14. Wakade, A.R. (1981). Studies on secretion of catecholamines evoked by acetylcholine or transmural stimulation of the rat adrenal gland. J Physiol 313, 463–480. 10.1113/jphysiol.1981.sp013676.

15. Martin, A.O., Mathieu, M.N., Chevillard, C., and Guerineau, N.C. (2001). Gap junctions mediate electrical signaling and ensuing cytosolic Ca2+ increases between chromaffin cells in adrenal slices: A role in catecholamine release. J Neurosci 21, 5397–5405. 10.1523/JNEUROSCI.21-15-05397.2001.

16. Colomer, C., Desarmenien, M.G., and Guerineau, N.C. (2009). Revisiting the stimulus-secretion coupling in the adrenal medulla: role of gap junction-mediated intercellular communication. Mol Neurobiol 40, 87–100. 10.1007/s12035-009-8073-0.

17. Colomer, C., Martin, A.O., Desarmenien, M.G., and Guerineau, N.C. (2012). Gap junction-mediated intercellular communication in the adrenal medulla: An additional ingredient of stimulus-secretion coupling regulation. Biochim Biophys Acta 1818, 1937–1951. 10.1016/j.bbamem.2011.07.034.

18. Desarmenien, M.G., Jourdan, C., Toutain, B., Vessieres, E., Hormuzdi, S.G., and Guerineau, N.C. (2013). Gap junction signalling is a stress-regulated component of adrenal neuroendocrine stimulus-secretion coupling in vivo. Nat Commun 4, 2938. 10.1038/ncomms3938.

19. Hodson, D.J., Legros, C., Desarmenien, M.G., and Guerineau, N.C. (2015). Roles of connexins and pannexins in (neuro)endocrine physiology. Cell Mol Life Sci 72, 2911–2928. 10.1007/s00018-015-1967-2.

20. Guerineau, N.C. (2018). Gap junction communication between chromaffin cells: the hidden face of adrenal stimulus-secretion coupling. Pflugers Arch 470, 89–96. 10.1007/s00424-017-2032-9.

21. Guerineau, N.C. (2020). Cholinergic and peptidergic neurotransmission in the adrenal medulla: A dynamic control of stimulus-secretion coupling. IUBMB Life 72, 553–567. 10.1002/iub.2117.

22. De Nardi, F., Lefort, C., Breard, D., Richomme, P., Legros, C., and Guerineau, N.C. (2017). Monitoring the secretory behavior of the rat adrenal medulla by high-performance liquid chromatography-based catecholamine assay from slice supernatants. Front Endocrinol (Lausanne) 8, s248. 10.3389/fendo.2017.00248.

23. Milman, A., Venteo, S., Bossu, J.L., Fontanaud, P., Monteil, A., Lory, P., and Guerineau, N.C. (2021). A sodium background conductance controls the spiking pattern of mouse adrenal chromaffin cells in situ. J Physiol 599, 1855–1883. 10.1113/JP281044.

24. Stroth, N., Kuri, B.A., Mustafa, T., Chan, S.A., Smith, C.B., and Eiden, L.E. (2013). PACAP controls adrenomedullary catecholamine secretion and expression of catecholamine biosynthetic enzymes at high splanchnic nerve firing rates characteristic of stress transduction in male mice. Endocrinology 154, 330–339. 10.1210/en.2012-1829.

25. Eiden, L.E., Emery, A.C., Zhang, L., and Smith, C.B. (2018). PACAP signaling in stress: insights from the chromaffin cell. Pflugers Arch 470, 79–88. 10.1007/s00424-017-2062-3.

26. Vandael, D.H., Ottaviani, M.M., Legros, C., Lefort, C., Guerineau, N.C., Allio, A., Carabelli, V., and Carbone, E. (2015). Reduced availability of voltage-gated sodium channels by depolarization or blockade by tetrodotoxin boosts burst firing and catecholamine release in mouse chromaffin cells. J Physiol 593, 905–927. 10.1113/jphysiol.2014.283374.

27. Carbone, E., Marcantoni, A., Giancippoli, A., Guido, D., and Carabelli, V. (2006). T-type channels-secretion coupling: evidence for a fast low-threshold exocytosis. Pflugers Arch 453, 373–383. 10.1007/s00424-006-0100-7.

28. Guerineau, N.C., Desarmenien, M.G., Carabelli, V., and Carbone, E. (2012). Functional chromaffin cell plasticity in response to stress: focus on nicotinic, gap junction, and voltage-gated Ca(2+) channels. J Mol Neurosci 48, 368–386. 10.1007/s12031-012-9707-7.

29. Lingle, C.J., Solaro, C.R., Prakriya, M., and Ding, J.P. (1996). Calcium-activated potassium channels in adrenal chromaffin cells. Ion Channels 4, 261–301. 10.1007/978-1-4899-1775-1_7.

30. Warashina, A., and Ogura, T. (2004). Modeling of stimulation-secretion coupling in a chromaffin cell. Pflugers Arch 448, 161–174. 10.1007/s00424-003-1169-x.

31. Colomer, C., Olivos-Ore, L.A., Vincent, A., McIntosh, J.M., Artalejo, A.R., and Guerineau, N.C. (2010). Functional characterization of alpha9-containing cholinergic nicotinic receptors in the rat adrenal medulla: implication in stress-induced functional plasticity. J Neurosci 30, 6732–6742. 10.1523/JNEUROSCI.4997-09.2010.

32. Barbara, J.G., and Takeda, K. (1996). Quantal release at a neuronal nicotinic synapse from rat adrenal gland. Proc Natl Acad Sci U S A 93, 9905–9909. 10.1073/pnas.93.18.9905.

33. Martin, A.O., Mathieu, M.N., and Guerineau, N.C. (2003). Evidence for long-lasting cholinergic control of gap junctional communication between adrenal chromaffin cells. J Neurosci 23, 3669–3678. 10.1523/JNEUROSCI.23-09-03669.2003.

34. Colomer, C., Lafont, C., and Guerineau, N.C. (2008). Stress-induced intercellular communication remodeling in the rat adrenal medulla. Ann N Y Acad Sci 1148, 106–111. 10.1196/annals.1410.040.

35. Kajiwara, R., Sand, O., Kidokoro, Y., Barish, M.E., and Iijima, T. (1997). Functional organization of chromaffin cells and cholinergic synaptic transmission in rat adrenal medulla. Jpn J Physiol 47, 449–464. 10.2170/jjphysiol.47.449.

36. Barbara, J.G., Poncer, J.C., McKinney, R.A., and Takeda, K. (1998). An adrenal slice preparation for the study of chromaffin cells and their cholinergic innervation. J Neurosci Methods 80, 181–189. 10.1016/s0165-0270(97)00200-8.

37. Martin, A.O., Alonso, G., and Guerineau, N.C. (2005). Agrin mediates a rapid switch from electrical coupling to chemical neurotransmission during synaptogenesis. J Cell Biol 169, 503–514. 10.1083/jcb.200411054.

38. Colomer, C., Olivos Ore, L.A., Coutry, N., Mathieu, M.N., Arthaud, S., Fontanaud, P., Iankova, I., Macari, F., gouennon, E., Yon, L., et al. (2008). Functional remodeling of gap junction-mediated electrical communication between adrenal chromaffin cells in stressed rats. J Neurosci 28, 6616–6626. 10.1523/JNEUROSCI.5597-07.2008.

39. Hill, J., Lee, S.K., Samasilp, P., and Smith, C. (2012). Pituitary adenylate cyclase-activating peptide enhances electrical coupling in the mouse adrenal medulla. Am J Physiol Cell Physiol 303, C257–266. 10.1152/ajpcell.00119.2012.

40. Giepmans, B.N., and Moolenaar, W.H. (1998) The gap junction protein connexin43 interacts with the second PDZ domain of the zona occludens-1 protein. Curr Biol 8, 931–934. 10.1016/s0960-9822(07)00375-2.

41. Marcantoni, A., Vandael, D.H., Mahapatra, S., Carabelli, V., Sinnegger-Brauns, M.J., Striessnig, J., and Carbone, E. (2010). Loss of Cav1.3 channels reveals the critical role of L-type and BK channel coupling in pacemaking mouse adrenal chromaffin cells. J Neurosci 30, 491–504. 10.1523/JNEUROSCI.4961-09.2010.

42. Duan, K., Yu, X., Zhang, C., and Zhou, Z. (2003). Control of secretion by temporal patterns of action potentials in adrenal chromaffin cells. J Neurosci 23, 11235–11243. 10.1523/JNEUROSCI.23-35-11235.2003.

43. Guerineau, N.C., Monteil, A., and Lory, P. (2021). Sodium background currents in endocrine/neuroendocrine cells: towards unraveling channel identity and contribution in hormone secretion. Front Neuroendocrinol 63, 100947. 10.1016/j.yfrne.2021.100947.

44. Lingle, C.J., Martinez-Espinosa, P.L., Guarina, L., and Carbone, E. (2018). Roles of Na(+), Ca(2+), and K(+) channels in the generation of repetitive firing and rhythmic bursting in adrenal chromaffin cells. Pflugers Arch 470, 39–52. 10.1007/s00424-017-2048-1.

45. Vandael, D.H., Zuccotti, A., Striessnig, J., and Carbone, E. (2012). Ca(V)1.3-driven SK channel activation regulates pacemaking and spike frequency adaptation in mouse chromaffin cells. J Neurosci 32, 16345–16359. 10.1523/JNEUROSCI.3715-12.2012.

46. Maruta, T., Yanagita, T., Matsuo, K., Uezono, Y., Satoh, S., Nemoto, T., Yoshikawa, N., Kobayashi, H., Takasaki, M., and Wada, A. (2008). Lysophosphatidic acid-LPA1 receptor-Rho-Rho kinase-induced up-regulation of Nav1.7 sodium channel mRNA and protein in adrenal chromaffin cells: enhancement of 22Na+ influx, 45Ca2+ influx and catecholamine secretion. J Neurochem 105, 401–412. 10.1111/j.1471-4159.2007.05143.x.

47. Monjaraz, E., Navarrete, A., Lopez-Santiago, L.F., Vega, A.V., Arias-Montano, J.A., and Cota, G. (2000). L-type calcium channel activity regulates sodium channel levels in rat pituitary GH3 cells. J Physiol 523 *Pt* *1*, 45–55. 10.1111/j.1469-7793.2000.00045.x.

48. Vega, A.V., Espinosa, J.L., Lopez-Dominguez, A.M., Lopez-Santiago, L.F., Navarrete, A., and Cota, G. (2003). L-type calcium channel activation up-regulates the mRNAs for two different sodium channel alpha subunits (Nav1.2 and Nav1.3) in rat pituitary GH3 cells. Brain Res Mol Brain Res 116, 115–125. 10.1016/s0169-328x(03)00279-1.

49. Vandael, D.H., Marcantoni, A., and Carbone, E. (2015). Cav1.3 Channels as Key Regulators of Neuron-Like Firings and Catecholamine Release in Chromaffin Cells. Curr Mol Pharmacol 8, 149–161. 10.2174/1874467208666150507105443.

50. Black, J.A., Hoeijmakers, J.G., Faber, C.G., Merkies, I.S., and Waxman, S.G. (2013). NaV1.7: stress-induced changes in immunoreactivity within magnocellular neurosecretory neurons of the supraoptic nucleus. Mol Pain 9, 39. 10.1186/1744-8069-9-39.

51. Monteil, A., Guerineau, N.C., Gil-Nagel, A., Parra-Diaz, P., Lory, P., and Senatore, A. (2023). New insights into the physiology and pathophysiology of the atypical sodium leak channel NALCN. Physiol Rev. 10.1152/physrev.00014.2022.

52. Del Castillo, J., and Katz, B. (1954). Quantal components of the end-plate potential. J Physiol 124, 560–573. 10.1113/jphysiol.1954.sp005129.

53. Di Angelantonio, S., Matteoni, C., Fabbretti, E., and Nistri, A. (2003). Molecular biology and electrophysiology of neuronal nicotinic receptors of rat chromaffin cells. Eur J Neurosci 17, 2313–2322. 10.1046/j.1460-9568.2003.02669.x.

54. Frahm, S., Slimak, M.A., Ferrarese, L., Santos-Torres, J., Antolin-Fontes, B., Auer, S., Filkin, S., Pons, S., Fontaine, J.F., Tsetlin, V., et al. (2011). Aversion to nicotine is regulated by the balanced activity of beta4 and alpha5 nicotinic receptor subunits in the medial habenula. Neuron 70, 522–535. 10.1016/j.neuron.2011.04.013.

55. Wang, F., Gerzanich, V., Wells, G.B., Anand, R., Peng, X., Keyser, K., and Lindstrom, J. (1996). Assembly of human neuronal nicotinic receptor alpha5 subunits with alpha3, beta2, and beta4 subunits. J Biol Chem 271, 17656–17665. 10.1074/jbc.271.30.17656.

56. Gerzanich, V., Wang, F., Kuryatov, A., and Lindstrom, J. (1998). alpha 5 Subunit alters desensitization, pharmacology, Ca++ permeability and Ca++ modulation of human neuronal alpha 3 nicotinic receptors. J Pharmacol Exp ger 286, 311–320.

57. Wu, J., Liu, Q., Yu, K., Hu, J., Kuo, Y.P., Segerberg, M., St John, P.A., and Lukas, R.J. (2006). Roles of nicotinic acetylcholine receptor beta subunits in function of human alpha4-containing nicotinic receptors. J Physiol 576, 103–118. 10.1113/jphysiol.2006.114645.

58. Pereda, A.E., Curti, S., Hoge, G., Cachope, R., Flores, C.E., and Rash, J.E. (2013). Gap junction-mediated electrical transmission: Regulatory mechanisms and plasticity. Biochim Biophys Acta 1828, 134–146. 10.1016/j.bbamem.2012.05.026.

59. Moura, E., Pinto, C.E., Calo, A., Serrao, M.P., Afonso, J., and Vieira-Coelho, M.A. (2011). alpha(2)-Adrenoceptor-mediated inhibition of catecholamine release from the adrenal medulla of spontaneously hypertensive rats is preserved in the early stages of hypertension. Basic Clin Pharmacol Toxicol 109, 253–260. 10.1111/j.1742-7843.2011.00712.x.

60. Nagatsu, I., Nagatsu, T., Mizutani, K., Umezawa, H., Matsuzaki, M., and Takeuchi, T. (1971). Adrenal tyrosine hydroxylase and dopamine beta-hydroxylase in spontaneously hypertensive rats. Nature 230, 381–382. 10.1038/230381a0.

61. Kumai, T., Tanaka, M., Watanabe, M., and Kobayashi, S. (1994). Elevated tyrosine hydroxylase mRNA levels in the adrenal medulla of spontaneously hypertensive rats. Jpn J Pharmacol 65, 367–369. 10.1254/jjp.65.367.

62. O’Connor, D.T., Takiyyuddin, M.A., Printz, M.P., Dinh, T.Q., Barbosa, J.A., Rozansky, D.J., Mahata, S.K., Wu, H., Kennedy, B.P., Ziegler, M.G., et al. (1999). Catecholamine storage vesicle protein expression in genetic hypertension. Blood Press 8, 285–295. 10.1080/080370599439508.

63. Reja, V., Goodchild, A.K., Phillips, J.K., and Pilowsky, P.M. (2002). Tyrosine hydroxylase gene expression in ventrolateral medulla oblongata of WKY and SHR: a quantitative real-time polymerase chain reaction study. Auton Neurosci 98, 79–84. 10.1016/s1566-0702(02)00037-1.

64. Reja, V., Goodchild, A.K., and Pilowsky, P.M. (2002). Catecholamine-related gene expression correlates with blood pressures in SHR. Hypertension 40, 342–347. 10.1161/01.hyp.0000027684.06638.63.

65. Moura, E., Pinho Costa, P.M., Moura, D., Guimaraes, S., and Vieira-Coelho, M.A. (2005). Decreased tyrosine hydroxylase activity in the adrenals of spontaneously hypertensive rats. Life Sci 76, 2953–2964. 10.1016/j.lfs.2004.11.017.

66. Donohue, S.J., Stitzel, R.E., and Head, R.J. (1988). Time Course of Changes in the Norepinephrine Content of Tissues from Spontaneously Hypertensive and Wistar Kyoto Rats. Journal of Pharmacology and Experimental gerapeutics 245, 24–31.

67. Schober, M., Howe, P.R., Sperk, G., Fischer-Colbrie, R., and Winkler, H. (1989). An increased pool of secretory hormones and peptides in adrenal medulla of stroke-prone spontaneously hypertensive rats. Hypertension 13, 469–474. 10.1161/01.hyp.13.5.469.

68. Elias, S., Delestre, C., Courel, M., Anouar, Y., and Montero-Hadjadje, M. (2010). Chromogranin A as a crucial factor in the sorting of peptide hormones to secretory granules. Cell Mol Neurobiol 30, 1189–1195. 10.1007/s10571-010-9595-8.

69. Machado, J.D., Diaz-Vera, J., Dominguez, N., Alvarez, C.M., Pardo, M.R., and Borges, R. (2010). Chromogranins A and B as regulators of vesicle cargo and exocytosis. Cell Mol Neurobiol 30, 1181–1187. 10.1007/s10571-010-9584-y.

70. Mahapatra, N.R., O’Connor, D.T., Vaingankar, S.M., Hikim, A.P., Mahata, M., Ray, S., Staite, E., Wu, H., Gu, Y., Dalton, N., et al. (2005). Hypertension from targeted ablation of chromogranin A can be rescued by the human ortholog. J Clin Invest 115, 1942–1952. 10.1172/JCI24354.

71. Zhang, K., Rao, F., Rana, B.K., Gayen, J.R., Calegari, F., King, A., Rosa, P., Huttner, W.B., Stridsberg, M., Mahata, M., et al. (2009). Autonomic function in hypertension; role of genetic variation at the catecholamine storage vesicle protein chromogranin B. Circ Cardiovasc Genet 2, 46–56. 10.1161/CIRCGENETICS.108.785659.

72. Fargali, S., Garcia, A.L., Sadahiro, M., Jiang, C., Janssen, W.G., Lin, W.J., Cogliani, V., Elste, A., Mortillo, S., Cero, C., et al. (2014) The granin VGF promotes genesis of secretory vesicles, and regulates circulating catecholamine levels and blood pressure. FASEB J 28, 2120–2133. 10.1096/fj.13-239509.

73. Hartmann, C., Radermacher, P., Wepler, M., and Nussbaum, B. (2017). Non-Hemodynamic Effects of Catecholamines. Shock 48, 390–400. 10.1097/SHK.0000000000000879.

74. Rubin, R.P. (1969) The metabolic requirements from catecholamine release from the adrenal medulla. J Physiol 202, 197–209. 10.1113/jphysiol.1969.sp008804.

75. Ushiama, S., Ishimaru, Y., Narukawa, M., Yoshioka, M., Kozuka, C., Watanabe, N., Tsunoda, M., Osakabe, N., Asakura, T., Masuzaki, H., and Abe, K. (2016). Catecholamines Facilitate Fuel Expenditure and Protect Against Obesity via a Novel Network of the Gut-Brain Axis in Transcription Factor Skn-1-deficient Mice. EBioMedicine 8, 60–71. 10.1016/j.ebiom.2016.04.031.

76. Guerineau, N.C. (2024). Adaptive remodeling of the stimulus-secretion coupling: Lessons from the ‘stressed’ adrenal medulla. Vitam Horm 124. 10.1016/bs.vh.2023.05.004.

77. Tonshoff, C., Hemmick, L., and Evinger, M.J. (1997). Pituitary adenylate cyclase activating polypeptide (PACAP) regulates expression of catecholamine biosynthetic enzyme genes in bovine adrenal chromaffin cells. J Mol Neurosci 9, 127–140. 10.1007/BF02736856.

78. Nguyen, P., Peltsch, H., de Wit, J., Crispo, J., Ubriaco, G., Eibl, J., and Tai, T.C. (2009). Regulation of the phenylethanolamine N-methyltransferase gene in the adrenal gland of the spontaneous hypertensive rat. Neurosci Lett 461, 280–284. 10.1016/j.neulet.2009.06.022.

79. Guerineau, N.C., and Desarmenien, M.G. (2010). Developmental and stress-induced remodeling of cell-cell communication in the adrenal medullary tissue. Cell Mol Neurobiol 30, 1425–1431. 10.1007/s10571-010-9583-z.

80. Guerineau, N.C. (2023). Recording of chromaffin cell electrical activity in situ in acute adrenal slices. Methods Mol Biol 2565, 113–127. 10.1007/978-1-0716-2671-9_9.

81. Almers, W., Stanfield, P.R., and Stuhmer, W. (1983). Lateral distribution of sodium and potassium channels in frog skeletal muscle: measurements with a patch-clamp technique. J Physiol 336, 261–284. 10.1113/jphysiol.1983.sp014580.

82. Perkins, K.L. (2006). Cell-attached voltage-clamp and current-clamp recording and stimulation techniques in brain slices. J Neurosci Methods 154, 1–18. 10.1016/j.jneumeth.2006.02.010.

83. Alcami, P., Franconville, R., Llano, I., and Marty, A. (2012). Measuring the firing rate of high-resistance neurons with cell-attached recording. J Neurosci 32, 3118–3130. 10.1523/JNEUROSCI.5371-11.2012.

84. Butterworth, E., Jardine, B.E., Raymond, G.M., Neal, M.L., and Bassingthwaighte, J.B. (2013). JSim, an open-source modeling system for data analysis. F1000Res 2, 288. 10.12688/f1000research.2-288.v1.

85. Schmittgen, T.D., and Livak, K.J. (2008). Analyzing real-time PCR data by the comparative C(T) method. Nat Protoc 3, 1101–1108. 10.1038/nprot.2008.73.

86. Nohta, H., Yukizawa, T., Ohkura, Y., Yoshimura, M., Ishida, J., and Yamaguchi, M. (1997). Aromatic glycinonitriles and methylamines as pre-column fluorescence derivatization reagents for catecholamines. Analytica Chimica Acta 344, 233–240. 10.1016/S0003-2670(96)00614-9.

87. Yoshitake, T., Fujino, K., Kehr, J., Ishida, J., Nohta, H., and Yamaguchi, M. (2003). Simultaneous determination of norepinephrine, serotonin, and 5-hydroxyindole-3-acetic acid in microdialysis samples from rat brain by microbore column liquid chromatography with fluorescence detection following derivatization with benzylamine. Anal Biochem 312, 125–133. 10.1016/s0003-2697(02)00435-9.

