## Supplementary information for "Adaptive remodeling of rat adrenomedullary stimulus-secretion coupling in response to a chronic hypertensive environment"

**Table S1: Passive membrane properties of WKY and SHR chromaffin cells.** Cells were recorded in the whole-cell configuration of the patch-clamp technique. The number of recorded cells is indicated in parentheses.

| | **WKY** | **SHR** | |
|---|---|---|---|
| Passive membrane properties | Mean ± SD | Mean ± SD | p value (unpaired t test) |
| Resting membrane potential, mV | -64.8 ± 6.4 (67) | -64.0 ± 7.4 (66) | 0.4951, ns |
| Input resistance, GΩ | 1.19 ± 0.36 (68) | 1.53 ± 0.42 (66) | <0.0001, **** |
| Membrane capacitance, pF | 8.94 ± 0.67 (104) | 8.74 ± 0.31 (96) | 0.009, ** |

**Table S2: Action potential properties of WKY and SHR chromaffin cells.** Depolarization-triggered and spontaneous action potentials were recorded in cells current-clamped at their resting potential. Evoked action potentials were triggered by a depolarizing step (+30 pA above the rheobase and 500 ms duration). For spontaneous APs, the analysis was performed in 30-50 serial APs/cell. All parameters were analyzed by Mini Analysis. The number of recorded cells is indicated in parentheses. *, p<0.05; **, p<0.01; ***, p<0.005, ****, p<0.001.

| | **WKY** (n = 25 cells) | **SHR** (n = 23 cells) | |
|---|---|---|---|
| **PROPERTIES OF EVOKED APs** | mean ± SD | mean ± SD | p value unpaired t test |
| rise time (ms) | 8.25 ± 0.17 | 8.42 ± 0.25 | 0.0086** |
| half-rise time (ms) | 1.18 ± 0.24 | 1.74 ± 0.66 | 0.0003*** |
| 10%-90% rise time (ms) | 3.41 ± 0.54 | 4.09 ± 0.83 | 0.0016** |
| 10%-90% slope (mV/ms) | 16.98 ± 4.67 | 13.44 ± 5.28 | 0.0175* |
| amplitude (mv) | 70.03 ± 7.38 | 62.56 ± 8.48 | 0.0021** |
| half width (ms) | 3.87 ± 1.01 | 5.73 ± 1.95 | 0.0001*** |
| decay time (ms) | 5.18 ± 1.74 | 7.99 ± 2.64 | p<0.0001**** |

| | **WKY** (n = 10 cells) | **SHR** (n = 8 cells) | |
|---|---|---|---|
| **PROPERTIES OF SPONTANEOUS APs** | mean ± SD | mean ± SD | p value Mann-Whitney test |
| rise time (ms) | 7.80 ± 0.32 | 8.23 ± 0.23 | 0.0085** |
| half-rise time (ms) | 0.64 ± 0.10 | 0.80 ± 0.10 | 0.0062** |
| 10%-90% rise time (ms) | 1.80 ± 0.33 | 2.50 ± 0.55 | 0.0014** |
| 10%-90% slope (mV/ms) | 41.95 ± 9.86 | 27.41 ± 6.61 | 0.0044** |
| amplitude (mv) | 89.51 ± 6.53 | 79.98 ± 7.78 | 0.0124* |
| half width (ms) | 3.32 ± 0.67 | 5.41 ± 2.16 | 0.0008*** |
| decay time (ms) | 6.00 ± 2.30 | 10.37 ± 6.60 | 0.0085** |

**Table S3: Primer sequences used for real-time RT-PCR**

| gene name | GenBank accession number | protein name | forward primer (5'-3') | reverse primer (5'-3') |
|---|---|---|---|---|
| *Scn1a* | NM_030875 | Nav1.1 | gttccgacatcgccagtt | catctcagtttcagtagttgttcca |
| *Scn2a* | NM_012647 | Nav1.2 | tggtgtccctggttggag | ccttatttctgtctcagtagttgtgc |
| *Scn3a* | NM_013119 | Nav1.3 | gcaccgtccattctaaccat | tttagcttcttgcataagaattgc |
| *Scn4a* | NM_013178 | Nav1.4 | ggcactgtctcgatttgagg | ttcatgatggaggggatagc |
| *Scn5a* | NM_013125 | Nav1.5 | tgccaccaatgccttgta | catgatgagcatgctaaagagc |
| *Scn8a* | NM_019266 | Nav1.6 | ggaagttttccatcatgaatcag | gctgttatgtcgggagagga |
| *Scn9a* | NM_133289 | Nav1.7 | cagcagatgttagaccgactca | actcgtgaactcagcagcag |
| *Scn1b* | NM_017288 | Nav1b | gcggagatggtgtactgctac | tcggaagtaatggccaggtat |
| *Scn2b* | NM_012877 | Nav2b | tggacttaccaggagtgtagca | ccagcttcaggttgatgatct |
| *Scn3b* | NM_139097 | Nav3b | ctgatacctttgcgagtcactg | catcatgatttccgagaccac |
| *Scn4b* | NM_001008880 | Navv4b | catcctgaagaagaccagagaga | agcagggcctcacactttt |
| *Cacna1c* | NM_012517 | Cav1.2 | tggctcacagaagtgcaaga | agcatttctgccgtgaaaag |
| *Cacna1d* | NM_017298 | Cav1.3 | ccatgctcactgtgttccag | ctcccatcctatcgcatcat |
| *Cacna1a* | NM_012918 | Cav2.1 | ctgctttgaagaggggacag | ctgacatttggtcccgttg |
| *Cacna1b* | NM_147141 | Cav2.2 | taagcgcatcaccgaatg | ggccaggacaatacagttgg |
| *Cacna1e* | NM_019294 | Cav2.3 | taccgcgcctggatagac | gctgatgttcccgagttttt |
| *Cacna1g* | NM_031601 | Cav3.1 | catctacttcattcttctcatcatcg | aactgcgtggcaatcacc |
| *Cacna1h* | NM_153814 | Cav3.2 | acggatactctgcagacagga | gctctgtgtagtctgggatgc |
| *Cacna1i* | NM_020084 | Cav3.3 | tctgcctcaatgttgtcacc | aagggctgtctctagggatgt |
| *Kcnma1* | NM_031828 | KCa1.1 | aagtaattccatcaagctggtga | ggtcccctgaattctccac |
| *Kcnn1* | NM_019313 | KCa2.1 | tcggaaacaccagcgtaagt | cttcacagtccggagcttct |
| *Kcnn2* | NM_019314 | KCa2.2 | tgcttgcttaccggaataatg | tagcttccttgccactacgg |
| *Kcnn3* | NM_019315 | KCa2.3 | cacgccaaagtcaggaaac | ccatcttgacacccctcagt |
| *Kcnn4* | NM_023021 | KCa3.1 | gcaagattgtctgcttgtgc | gccaccacagccaatagtaga |
| *Kcnt1* | NM_021853 | KCa4.1 | ggtcaagaaccgaatgaagc | tggctgttaaatttgctacgtc |

| | | | | |
|---|---|---|---|---|
| *Kcnt2* | NM_198762 | KCa4.2 | ccagcaaatatgagattcatgc | acctcgttctcgctctttctt |
| *Chrna3* | NM_052805 | α3nAChR | tccatgctgatgctggtg | tcttcgaacaggtactggaaca |
| *Chrna4* | NM_024354 | α4nAChR | ccagtacattgcagaccacct | gccacgtatttccagtcctc |
| *Chrna5* | NM_017078 | α5nAChR | cacgtcgtgaaagagaacga | ccacaaaaacatccgatcaa |
| *Chrna7* | NM_012832 | α7nAChR | ggcaaaatgcctaagtggac | cttcatgcgcagaaaccat |
| *Chrnb2* | NM_019297 | ß2nAChR | caactcaatggcgctgttc | cactagccgctcctctgtgt |
| *Chrnb4* | NM_052806 | ß4nAChR | tcctctcagctcatctccatc | tgatctgttctcgctcattca |
| *Gjd2* | NM_019281 | Cx36 | cggtactgcccagtctttgt | caccaccacagtcaacagga |
| *Gja1* | NM_012567 | Cx43 | cccgacgacaaccagaatg | tggctaatggctggagttc |
| *Tjp1* | NM_001106266 | ZO-1 | tcaacacatgaagatgggattc | cgcaaacccacactatctcc |
| *Gusb* | NM_017015 | Gus | ctctggtggccttacctgat | cagactcaggtgttgtcatcg |
| *Hprt* | NM_012583 | Hprt | gaccggttctgtcatgtcg | acctggttcatcatcactaatcac |
| *Gapdh* | NM_017008 | Gapdh | tgggaagctggtcatcaac | gcatcaccccatttgatgtt |

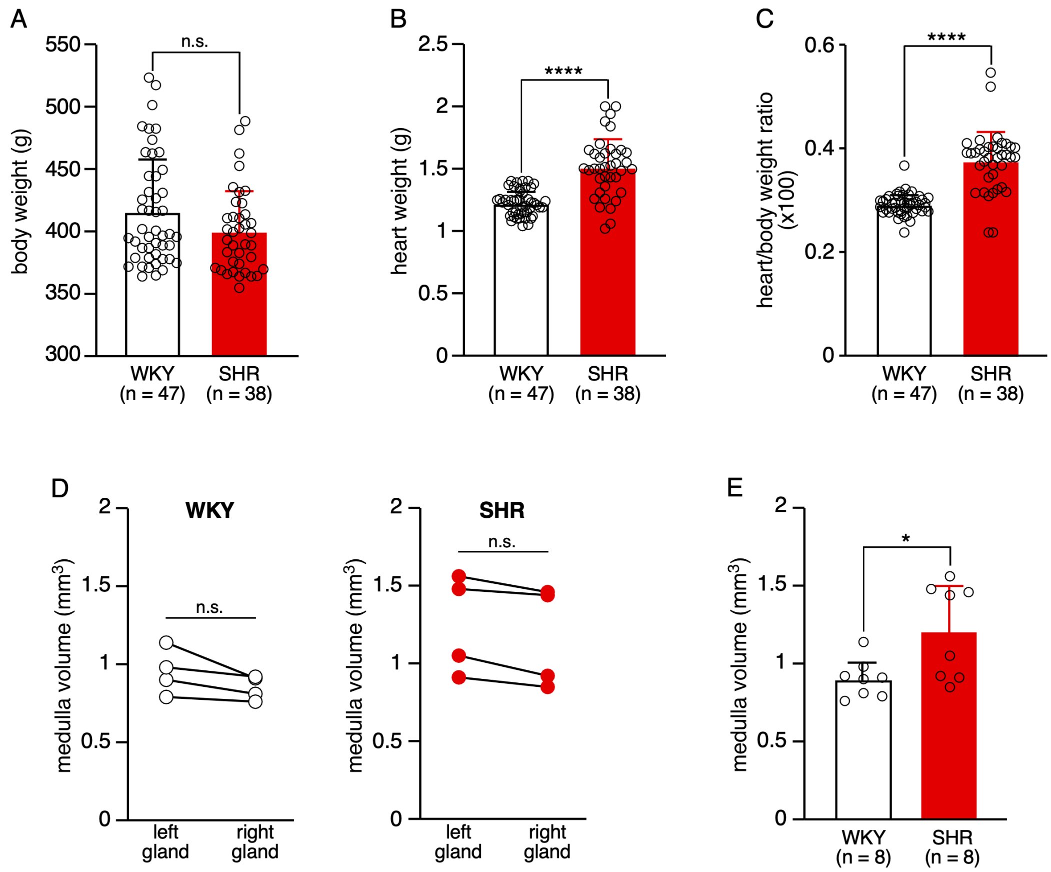

**Figure S1: Validation of phenotypic traits of SHRs compared to WKY rats. A.** No overt difference in body weight in the two rat strains (401.4 ± 33.3 g, n = 38) and WKY rats (417.2 ± 43.6 g, n = 47, p = 0.0689, unpaired t test). **B.** Significant increase in heart weight in hypertensive animals compared to normotensive animals (1.51 ± 0.23 g, n = 38 *versus* 1.22 ± 0.10 g, n = 47 in WKY rats, p<0.0001, unpaired t test). **C.** Enhanced heart/body weight ratio in SHRs (0.38 ± 0.06, n = 38 *versus* 0.29 ± 0.02, n = 47 in WKY rats, p<0.0001, unpaired t test). **D-E.** Comparison of the adrenal medulla volume between WKY rats and SHRs. The medulla volume was calculated from acute slices (see Material and Methods). **D.** No difference between the left and the right adrenals (0.85 ± 0.08 $mm^3$, n = 4 and 0.95 ± 0.15 $mm^3$, n = 4 for the right and left WKY glands, respectively, p = 0.125, and 1.17 ± 0.33 $mm^3$, n = 4 and 1.25 ± 0.32 $mm^3$, n = 4 for the right and left SHR glands, respectively, p = 0.125, Wilcoxon matched-pairs signed-rank test) **E.** Significantly greater medulla volume in SHRs *versus* WKY rats. (1.21 ± 0.30 $mm^3$, n = 8 glands for SHRs *versus* 0.90 ± 0.12 $mm^3$, n = 8 glands for WKY rats, p = 0.0351, Mann-Whitney test). For each rat, the volume represents the average volume of the two glands.

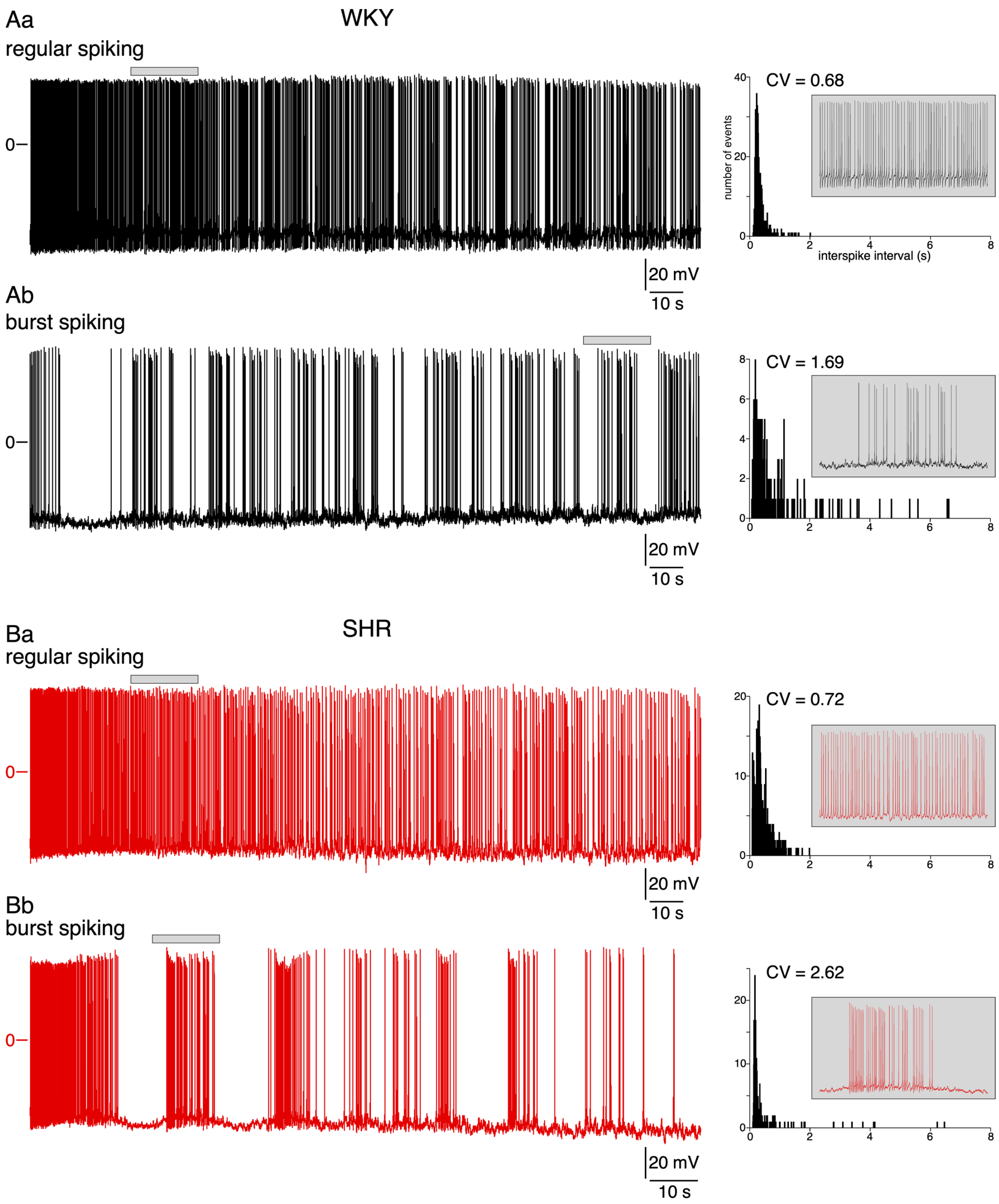

**Figure S2: Whole-cell recordings of spontaneously spiking chromaffin cells in adrenal acute slices: occurrence of both regular and burst firing.** Action potentials were recorded at resting membrane potential (no current injection), in normotensive (**A**) and hypertensive (**B**) rats. **A.** Representative chart recordings of two cells exhibiting a regular firing (**Aa**) or a burst firing (**Ab**). The histograms on the right illustrate the distribution of the inter-spike intervals (10 ms bin), from which the coefficients of variation (CV) were calculated. Insets: expanded time scale illustrating a 20-s spiking period. **B.** Same recording conditions and analysis performed in two SHR chromaffin cells.

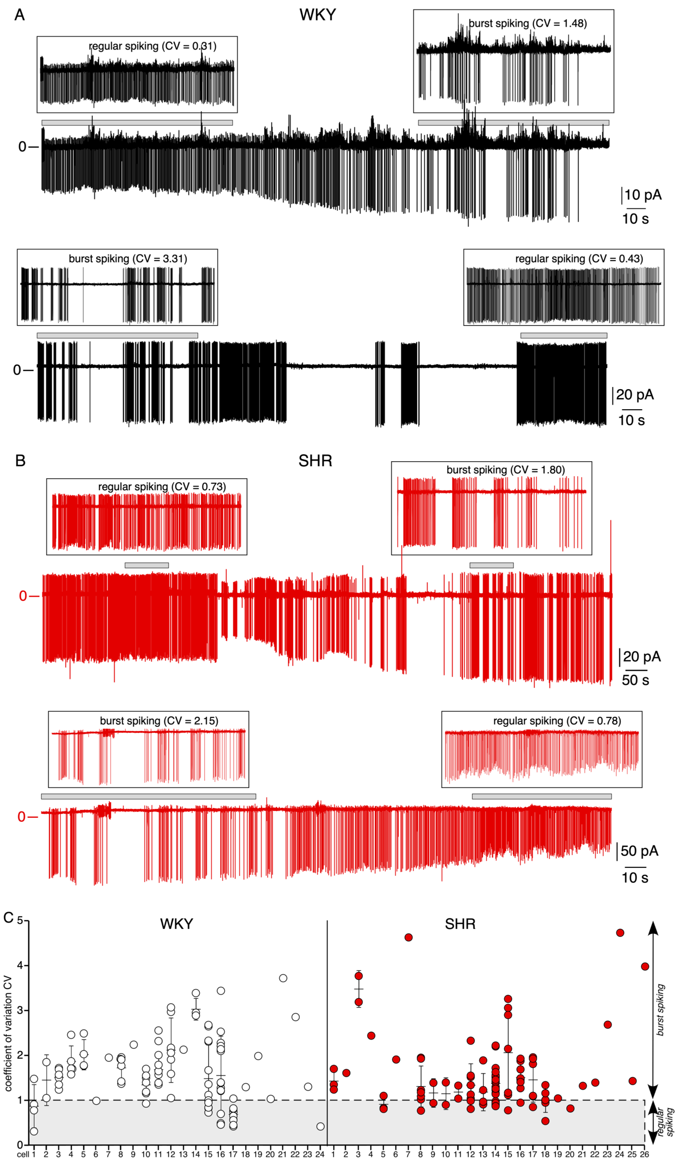

**Figure S3: Dynamic switch between regular and bursting spiking patterns in a same chromaffin cell, in WKY rats and SHRs. A.** Example of two WKY cells whose spontaneous electrical activity was recorded for 4-5 minutes and in which the spiking pattern switched from a regular to a bursting mode or vice versa. **B.** Same recordings in two SHR chromaffin cells, in which spontaneous action potentials were monitored for 4-20 min. Highlighted sections in A and B: expanded time scale (100 s) showing representative periods during which cells fired either regularly or in bursts. For each 100 s recording period, the distributions of inter-spike intervals were plotted and the associated CV calculated, as indicated in Figure S2. **C.** Distribution of CV values for each individual cell. All CV values were calculated from a 100 s recording (1-15 runs/cell in WKY rats and 1-21 runs/cell in SHRs). Note the wide distribution, both in normotensive and hypertensive animals, not only between cells but also between series of a same cell.

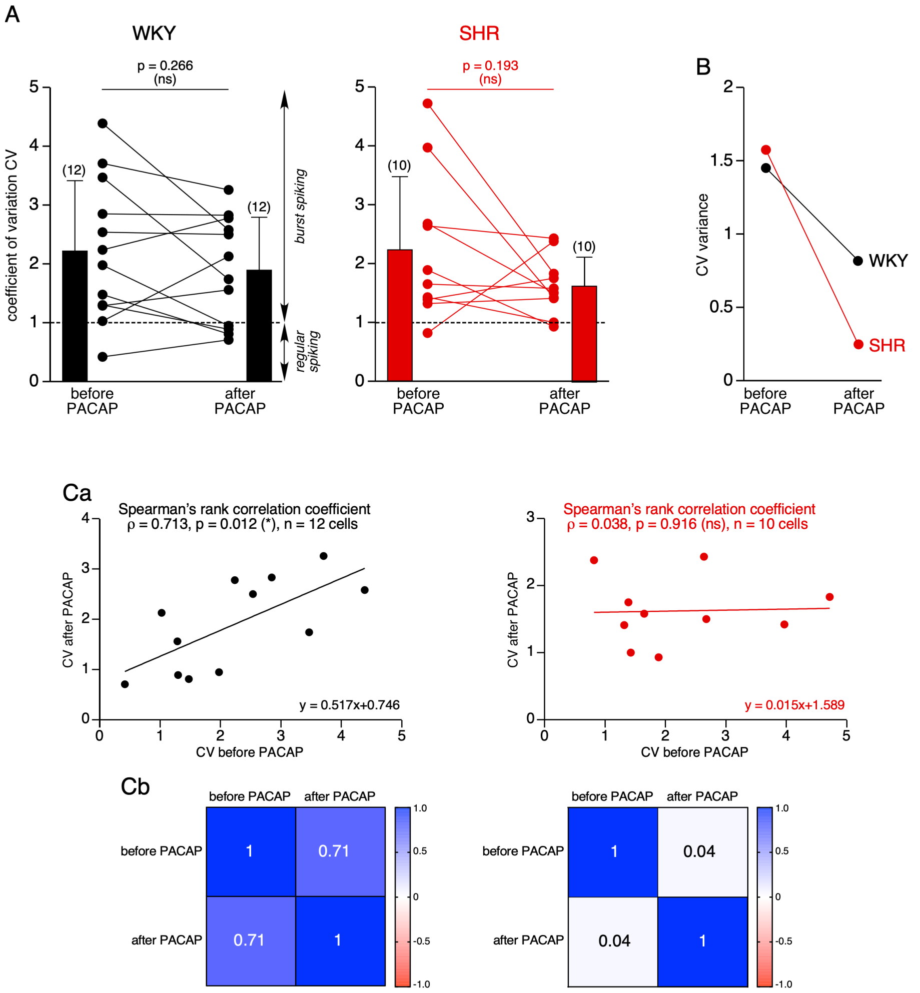

**Figure S4: Analysis of CV changes in response to PACAP.** Chromaffin cells from WKY rats and SHRs were exposed to 0.1-10 µM PACAP. Basal and PACAP-stimulated electrical activity were recorded in the loose cell-attached configuration (clamped at 0 mV). **A.** For each cell (12 WKY cells and 10 SHR cells), a CV was calculated for the basal condition (before PACAP) and for PACAP-exposed condition (after PACAP). Although CV tends to decrease in response to PACAP, the result is not significant, neither in normotensive or hypertensive animals (2.22 ± 1.20 before PACAP and 1.89 ± 0.91 after PACAP, n = 12 cells for WKY rats, p = 0.2661, Wilcoxon matched-pairs signed-rank test and 2.25 ± 1.26 before PACAP and 1.62 ± 0.50 after PACAP, n = 10 cells, p = 0.1934, Wilcoxon matched-pairs signed-rank test). **B.** Reduced CV variance after PACAP exposure, in particular in SHRs, indicating that PACAP acts by homogenizing the firing pattern. **C.** Loss of the correlation between CV after PACAP and CV before PACAP in SHRs (**Ca**) (ρ = 0.713 in WKY rats, p = 0.0012, n = 12 cells, Spearman's rank correlation coefficient *versus* ρ = 0.038 in SHRs, p = 0.916, n = 10 cells, Spearman's rank correlation coefficient) and associated table of correlation (**Cb**).

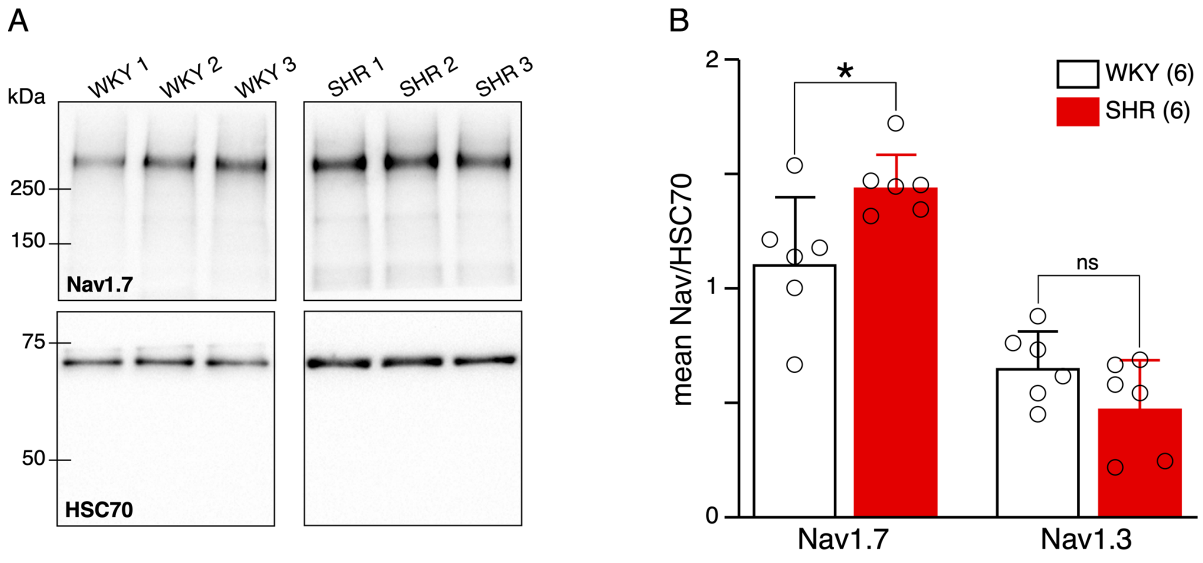

**Figure S5: Increased expression of Na$_V$1.7 protein in SHRs.** The adrenomedullary tissue was macrodissected in 6 WKY rats and 6 SHRs. **A.** Western blot illustration of Na$_V$1.7 in 3 WKY rats and 3 SHRs. Intensities of Na$_V$1.7 bands were normalized to those of HSC70. **B.** Pooled data showing that Na$_V$1.7 protein expression is significantly upregulated in SHRs (Na$_V$1.7/HSC70 ratio: 1.11 ± 0.28 for WKY rats, n = 6 and 1.45 ± 0.14 for SHRs, n = 6, p = 0.0411, Mann-Whitney test). By contrast, the expression of Na$_V$1.3 protein, the more expressed protein of the Nav family in rat chromaffin cells, is not modified in hypertensive rats compared to normotensive rats.

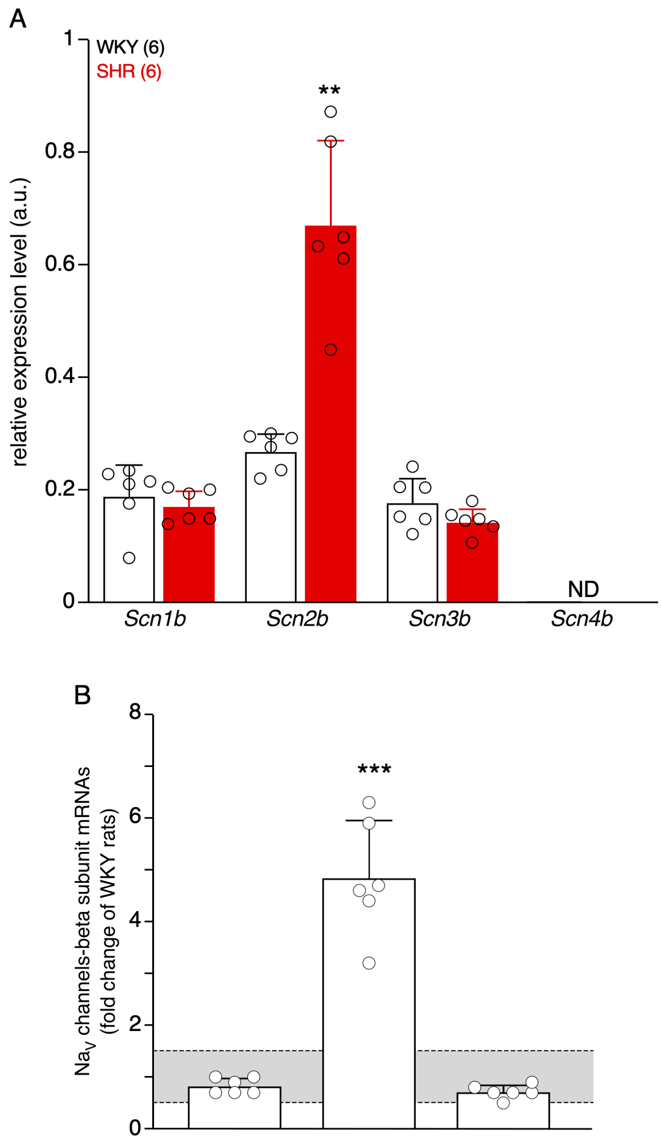

**Figure S6: Expression changes in transcripts encoding auxiliary β subunits of voltage-gated $Na^+$ channels in SHRs.** Changes in mRNA expression levels were assessed by real-time RT-PCR in macrodissected adrenal medullary tissues from 6 WKY rats and 6 SHRs. **A.** Of the four existing β subunits, only transcripts encoding β1 (*Scn1b*), β2 (*Scn2b*) and β3 (*Scn3b*) were detected, with a significant increase for *Scn2b* (relative expression of 0.27 ± 0.03 for WKY rats, n = 6 and 0.67 ± 015 for SHRs, n = 6, p = 0.0022, Mann-Whitney test). The *Scn4b* mRNA encoding β4 subunit was not detected (ND). **B.** Fold changes in SHRs, as compared to WKY rat. A significant change occurs for the gene encoding $Na_V$β2 subunit (4.8-fold). Fold change values were determined according to Livak's method (see Material and Methods). The Shapiro-Wilk test was used to analyze the normality of data distribution, and parametric or non-parametric unpaired tests were used when appropriate. Fold changes between x0.5 and x1.5 (grey area) are considered irrelevant.

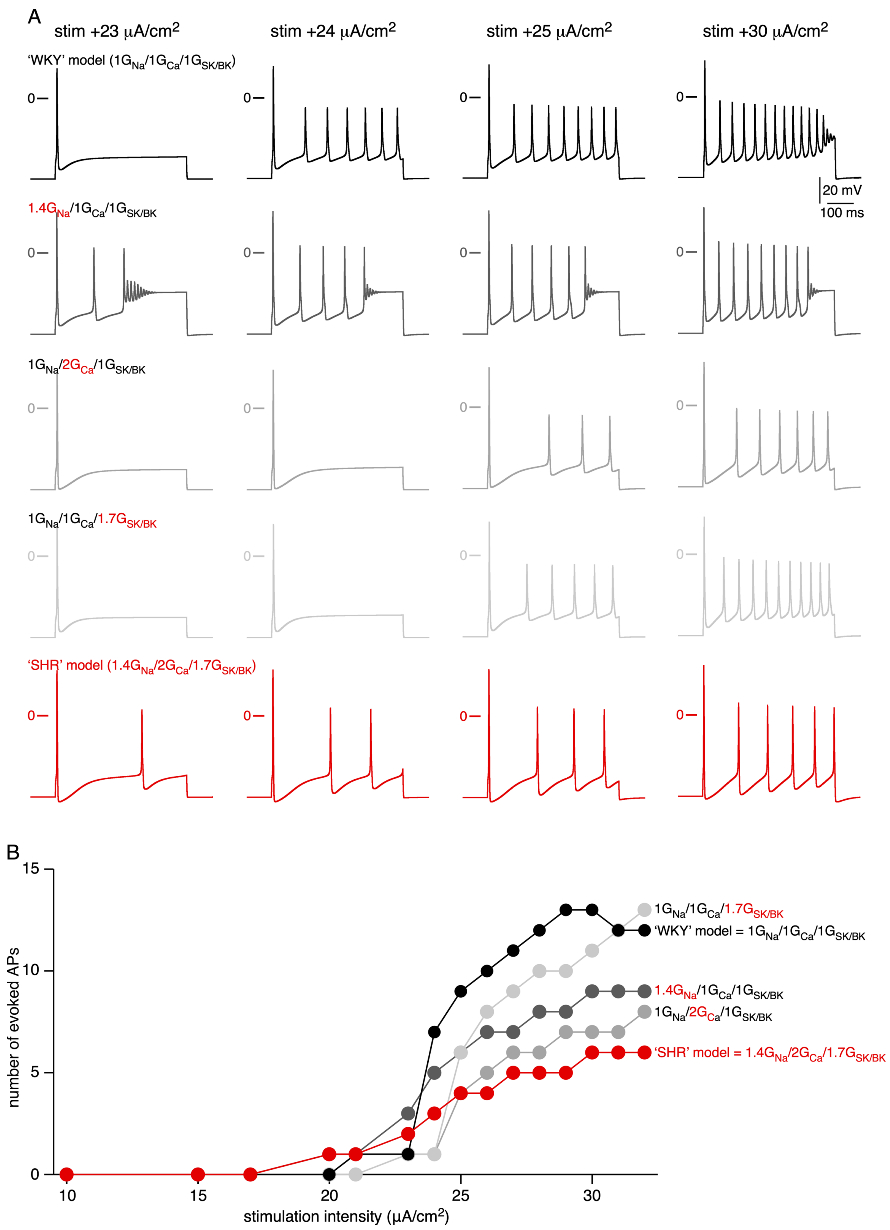

**Figure S7: Impact of changes in $G_{Na}$, $G_{Ca}$ and $G_{SK}$ conductances on the electrical firing of computed rat chromaffin cells.** 'WKY' and 'SHR' models were built from the rat chromaffin cell numerical simulation developed by Warashina and Ogura. **A.** Electrical firing evoked at four stimulation intensities (from 23 to 30 µA/cm$^2$, 500 ms duration). The 'SHR'-like condition (red traces) corresponds to 1.4$G_{Na}$, 2$G_{Ca}$ and 1.7$G_{SK/BK}$ 'WKY' model (see Material and Methods). In the intermediate conditions (light, medium and dark grey traces), either $G_{Na}$, $G_{Ca}$ or $G_{SK/BK}$ were modified by their respective multiplication coefficients, leaving the two other conductance unchanged. **B.** Pooled data showing that the most pronounced reduction in AP frequency is found when $G_{Na}$, $G_{Ca}$ and $G_{SK/BK}$ are modified simultaneously, particularly at high stimulation intensities.
